# Chromosome territory reorganization through artificial chromosome fusion is negligible to cell fate determination and mouse development

**DOI:** 10.1101/2022.10.09.511475

**Authors:** Yuang Wang, Zhen Qu, Yi Fang, Yulong Chen, Jiayin Peng, Jiawen Song, Jinsong Li, Jiantao Shi, Jin-Qiu Zhou, Yun Zhao

## Abstract

**Summary:** Chromosomes occupy discrete spaces in the interphase cell nucleus, called chromosome territory. The structural and functional relevance of chromosome territory remains elusive. We fused chromosome 15 and 17 in mouse haploid embryonic stem cells (haESCs), resulting in distinct changes of territories in the corresponding native chromosomes, but affecting little on gene expression, pluripotency and gamete functions of haESCs. The karyotype engineered haESCs were successfully implemented in cultivating heterozygous (2n=39) and homozygous (2n=38) mouse models. The mice containing fusion-chromosome are fertile, and their representative tissues and organs display no phenotypic abnormalities, suggesting mouse development unscathed. These results indicate that the mammalian chromosome architectures are highly resilient, and reorganization of chromosome territories can be readily tolerated during cell differentiation and mouse development.

## Introduction

Chromosome is the main carrier of genetic information, and is responsible for the transmission of genetic materials from parents to offspring. The number of chromosomes found in natural eukaryotic species ranges from one to thousands (Crosland and Crozier, 1986; Khandelwal, 1990). In most eukaryotic cells, chromosomes appear as chromatin during the interphase of the cell cycle and as linear “rods” during the mitotic phase (Bolzer et al., 2005). For a chromosome to function stably and effectively across generations, it must have a centromere and two telomeres. Centromere is a specific locus on a chromosome for assembly of the kinetochore, which is responsible for microtubule attachment and precise chromosome segregation during cell division (Cheeseman and Desai, 2008), while telomeres are the physical ends of a chromosome that protect chromosome from degradation (Bianchi and Shore, 2008; O’Sullivan and Karlseder, 2010).

Chromosomes do not seem to be randomly distributed or intermingled with each other in the nucleus. Instead, each chromosome occupies a distinct spatial volume in the interphase nucleus, called chromosome territory (Cremer and Cremer, 2001; Cremer and Cremer, 2010; Lamond and Earnshaw, 1998; Meaburn and Misteli, 2007). In addition, the territory of a particular chromosome may be different in different cell types (Cremer et al., 2001; Croft et al., 1999; Meaburn and Misteli, 2007). Accordingly, a proximal positioning of adjacent chromosomes may also be meaningful, e.g. regulating chromatin activities (Branco and Pombo, 2006a; Stevens et al., 2017; Tanabe et al., 2002). But what determines chromosome territory and how it regulates genome function are unclear.

The chromosome number in naturally evolved house mice *Mus musculus domesticus*, which have populated in Western Europe and North Africa, ranges from 2n=40 to 2n=22 (Capanna and Redi, 1995; Nachman and Searle, 1995; PiáLek et al., 2005). Some of their chromosomes are metacentric, i.e. the centromere is at the middle of each chromosome due to fusions of two telocentric chromosomes, which are commonly found in the laboratory mouse (e.g. C57BL/6) (Garagna et al., 2014). In addition, Muntjac deer (Muntiacus, Muntiacinae, Cervidae) have evolved quite diverse karyotypes (e.g. 2n = 46 of *M. reevesi* and 2n = 6/7 of *M. muntjak vaginalis*) through chromosome translocation, tandem fusion, and pericentric inversion (Fontana and Rubini, 1990; Neitzel, 1987; Yin et al., 2021). Recently, deliberate artificial chromosome engineering has succeeded in generating single-chromosomal *Saccharomyces cerevisiae* and *Schizosaccharomyces pombe* strains, which shows drastic changes in global chromosome structures, but grow as robustly as the naturally evolved strains (Gu et al., 2022; Luo et al., 2018; Shao et al., 2018). These lines of evidence suggest that chromosome architecture in eukaryotes is highly resilient, and chromosome territories could be self-organizing representations of the genome, or simply be a manifestation of random chromatin collisions driven by intrinsic interactions between chromatin loci and/or geometric constraints within the nucleus.

In order to experimentally address whether the high plasticity of chromosome architecture is a ubiquitous characteristic of eukaryotic genomes, we employed haploid embryonic stem cells (haESCs) and mouse models to test the effects of extreme chromosome territory changes on stem cell pluripotency, cell fate determination and mouse development.

## Results

### Construction of chromosome fusion haploid embryonic stem cells

We used CRISPR-Cas9 to induce double-strand-breaks in chromosome 15 and 17 in mouse haESCs, namely H19ΔDMR-IGΔDMR-AGH (hereafter referred as WT) (Zhong et al., 2015). The guide RNA (gRNA) targeting sites were in the regions of distal telomere (D-telomere) region of Chr15 and sub-centromeric telomere (C-telomere) region of Chr17, respectively (Figure 1A and Figure S1A). The broken chromosomes might fuse together by non-homologous-end joining (NHEJ) (Figure 1A), an active DNA repair mechanism intrinsic to cells (Sharma and Raghavan, 2016).

**Figure 1.**
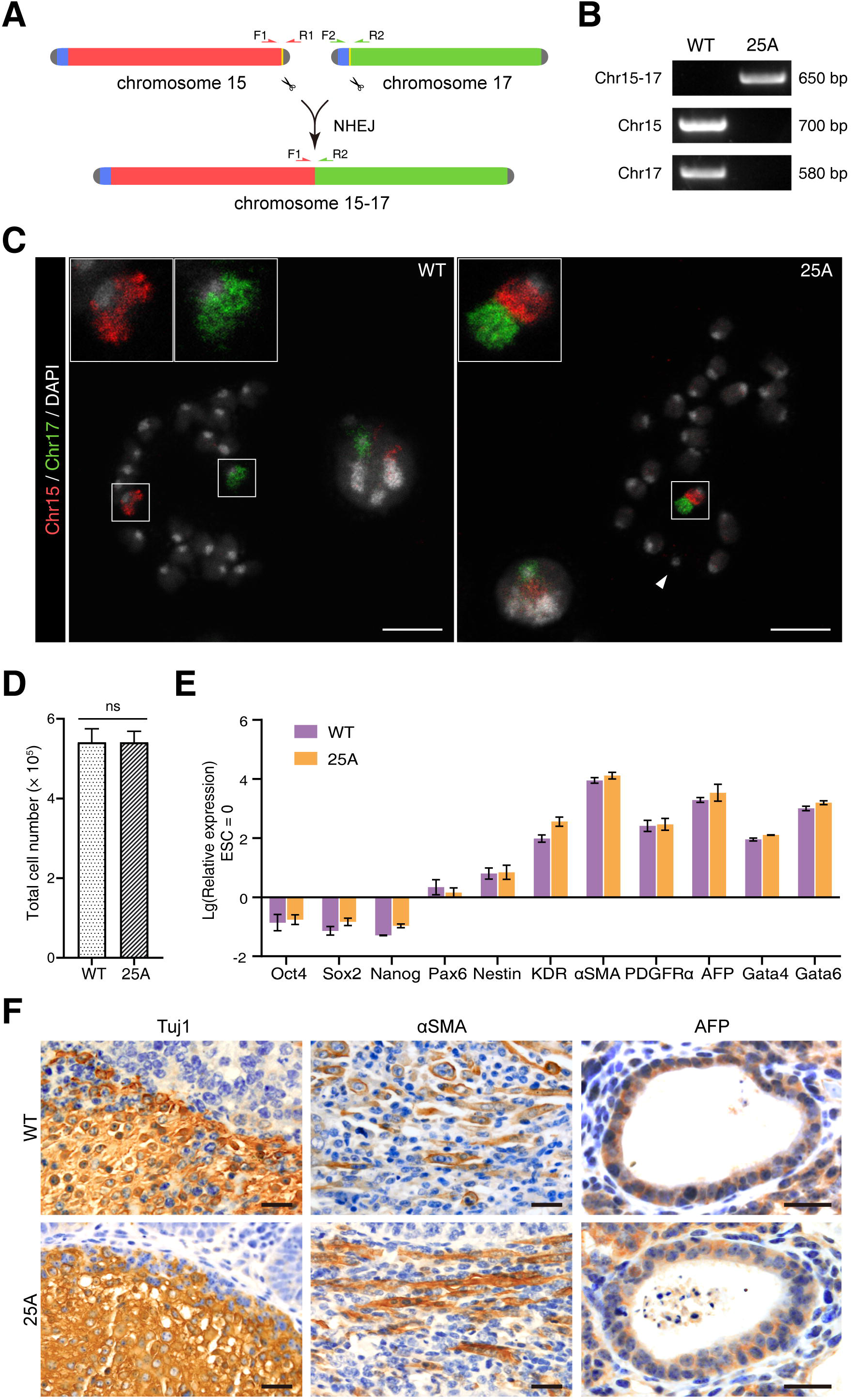
CRISPR/Cas9 mediated site-specific chromosome breaks and Chr15-17 fusion in haESCs. (A) Schematic showing the experimental strategy for generating site-specific chromosome fusion of Chr15 (red) and Chr17 (green) in haESCs. Two sgRNAs guide Cas9 (scissors) to the indicated target sites (yellow) located near distal telomere region (D-telomere, grey) of Chr15 and sub-centromeric telomere region (C-telomere, blue) of Chr17, respectively. Chromosome fusion occurred between two target sites is detected by cross-chromosomal PCR. Chr15 without D-telomere and Chr17 without C-telomere are ligated through NHEJ pathway, generating Chr15-17 fusion. Primers are designed at the upstream and downstream of each sgRNA target site. (B) PCR analysis of 25A haESCs with primers mentioned in (A). Cross-chromosomal PCR with primer pairs ‘F1’ and ‘R2’ amplified a ~ 650 bp band only in 25A, while ‘F1’ and ‘R1’ on Chr15 and ‘F2’ and ‘R2’ on Chr17 amplified a ~700 bp and a ~580 bp band, respectively only in WT but not in 25A. (C) Fluorescent images of the metaphase chromosomes of WT and 25A haESCs labelled with whole painting probes of Chr15 (red) and Chr17 (green). Insets zoomed-in showing the Chr15 and Chr17 in WT and the fused Chr15-17 in 25A. The mini-chromosome is indicated with white arrowhead. Scale bar: 10 μm. (D) Proliferation rates examined by total cell number of WT and 25A haESCs. ns: not significant. (E) Real time PCR analysis of the expression levels of pluripotency marker genes (*Oct4, Sox2* and *Nanog*) and differentiation-related genes (Ectoderm: *Pax6, Nestin*; Mesoderm: *KDR*, α*SMA, PDGFR*α; Endoderm: *AFP, Gata4, Gata6*) in differentiated WT and 25A cells. The expression levels were Lg transformed. Data are represented as the mean ± SD, n = 3. Gene expression levels were not significant between WT and 25A cells. (F) Paraffin sections of teratomas formed by WT and 25A cells were stained with three germ-layer markers including the ectoderm marker Tuj1, the mesoderm marker αSMA and the endoderm marker AFP. Scale bar: 20 μm.

The haESCs were transfected with CRISPR-Cas9 vectors, and potential clones were screened by cross-chromosomal PCR using primer pairs ~400 bp upstream and ~250 bp downstream of the respective gRNA targeting sites in Chr15 and Chr17 (Figure 1A). Two clones namely 25A and 42E showed an amplified band with the expected length (Figure 1B and Figure S1B). Further sequencing results confirmed that both PCR products matched the sequences adjacent to the respective gRNA targeting sites in Chr15 and Chr17 (Figure S1C), with 9 bp and 7 bp deletion at the junction sites, respectively, indicating that chromosome fusion in both clones is likely mediated by NHEJ.

Next, we performed chromosome FISH analysis by employing whole painting probes to verify chromosome fusion at the cellular level. In WT haESCs, the Chr15 and Chr17 were labelled with red and green fluorescent probes, respectively. In 25A and 42E cells, one half of a chromosome was labeled with red fluorescence, while the other half was labeled with green fluorescence, indicating the fusion of Chr15 and Chr17 (Figure 1C and Figure S1D). Notably, in either 25A or 42E cell line, there was a mini-chromosome (indicated by white arrowhead in Figure 1C and Figure S1D) that was not seen in the WT haESCs. We speculated that this mini-chromosome was the abandoned C-telomere of Chr17, and was stably maintained during the passages of the haESCs. Consistently, karyotype analysis also showed that Chr15 and Chr17 were fused (red arrowhead in Figure S1E, S1F), and the mini-chromosome was retained in 25A and 42E cells (blue arrowhead in Figure S1E, S1F).

### Chromosome fusion in haESCs does not affect cell pluripotency

To address whether chromosome fusion in haESCs affects cellular functions, we examined cell morphology, proliferation rate, karyotype stability and differentiation potentials. We isolated the haploid (G0/G1 phase) 25A cells by FACS (Figure S1G), and there was no significant difference between 25A and WT in terms of proliferation rate and colony morphology (Figure 1D and Figure S1H), and the karyotype of 25A was stably maintained after 25 passages (Figure S1I, S1J), suggesting that chromosome fusion does not affect mitosis.

Chromosome fusion 25A cells expressed pluripotency marker genes, including *Oct4, Sox2* and *Nanog*, which were not significantly different from WT (Figure S1K). To assess whether 25A cells remain pluripotent, we induced *in vitro* differentiation by removing leukemia inhibitory factor (LIF) and two differentiation inhibitors (CHIR99021, PD0325901) in the culture medium (Figure S1L). The cells showed differentiation morphology after two weeks (Figure S1M), in coincidence with the down-regulation of pluripotency marker genes and the up-regulation of the three germ layers’ differentiation related genes (Ectoderm marker genes: *Pax6, Nestin*; Mesoderm marker genes: *KDR*, α*SMA, PDGFR*α; Endoderm marker genes: *AFP, Gata4, Gata6*) (Figure 1E). Furthermore, subcutaneous injection of 25A cells into immunodeficient mice resulted in the formation of teratomas, which contained three germ layers identified by immunohistochemical (IHC) staining (Figure 1F). Collectively, these results indicate that chromosome fusion in haESCs does not affect cell pluripotency.

### Chromosome fusion leads to rearrangement of chromosome territory

There have been indications that different chromosomes occupy different spaces in cell nucleus, and adjacent positioning of chromosomes suggests their interactions are significant (Meaburn and Misteli, 2007). To explore the effect of chromosome fusion on chromosome territories, we performed whole-genome chromosome conformation analysis on diploid 25A and WT cells with high-coverage Hi-C sequencing (~100 ×). In the genome-wide contact matrixes, the inter-chromosomal interactions of Chr15 and Chr17 were largely random in WT cells (Figure 2A, Figure S2A, S2C, S2E), while significantly enhanced in 25A cells presumably because the chromosome fusion linked two chromosomes, resulting in the significant emergence of new intra-chromosome interactions (Figure 2A, Figure S2B, S2D, S2F). The distribution of contact probabilities as a function of genomic distances in fused Chr15-17 is indistinguishable from that of a single chromosome (e.g. Chr1 in WT or 25A) (Figure 2B), indicating that chromosome fusion has not only disrupted the original territories of native Chr15 and 17, but also established a new territory in the fused chromosome.

**Figure 2.**
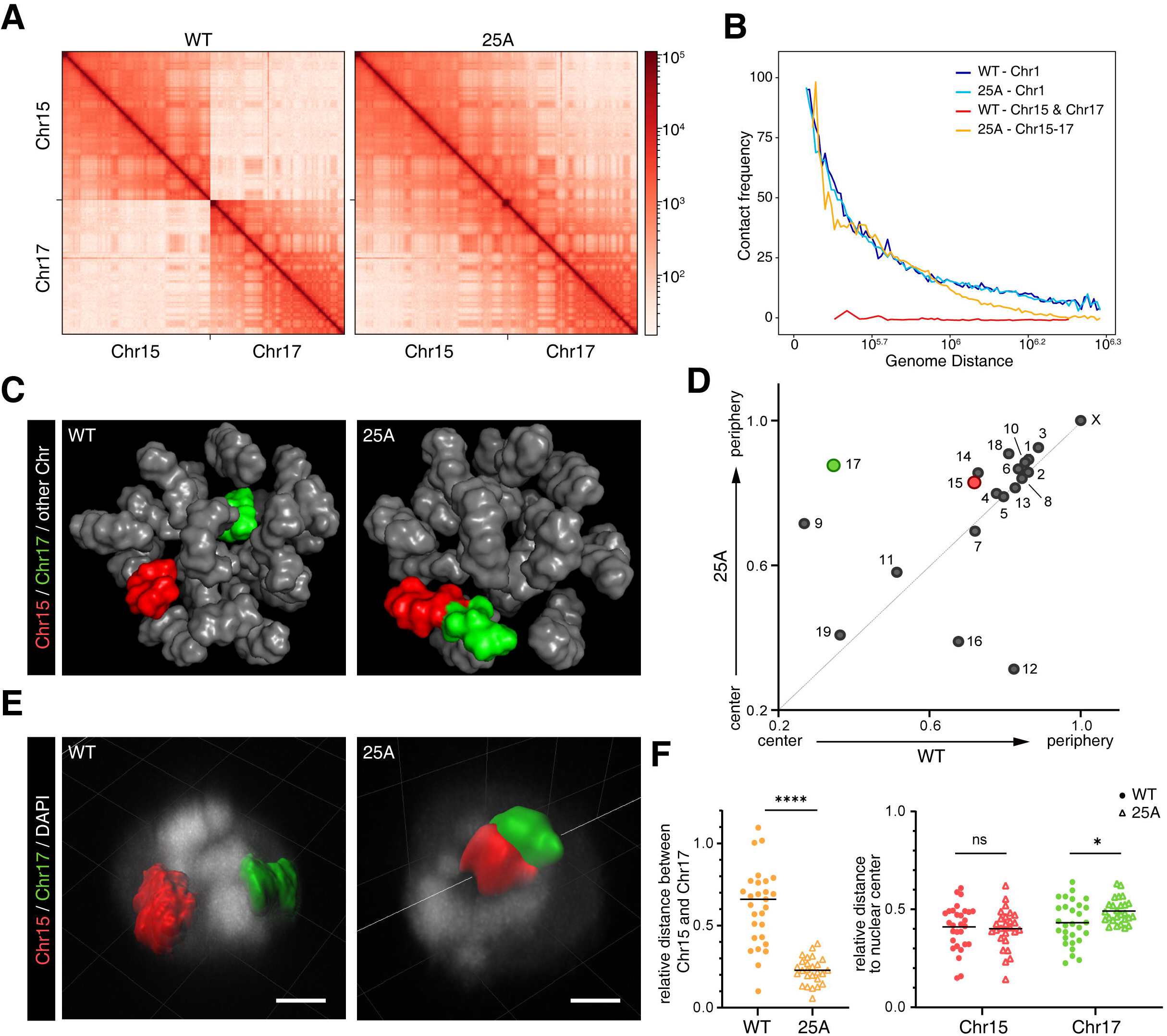
Change of chromosome territory after chromosome 15 and 17 fusion. (A)Contact matrixes at 1Mb resolution on Chr15 and 17 of WT and 25A cells. (B) Contact frequency curves showing contact probabilities as a function of genomic distances around fusion site (within 1 Mb) on Chr15 and Chr17. Chr1 was included as a single chromosome control. (C) Inferred 3D genome structures in WT and 25A cells using Hi-C data. Chr15 was shown in red and Chr17 in green. (D) Radial distributions of each chromosome relative to nucleus center in WT (horizontal axis) and 25A (vertical axis) derived from inferred 3D models (C). (E) 3D-FISH reconstruction of the Chr15 and Chr17 in WT and 25A haESCs. Chr15 was labeled in red and Chr17 in green. Scale bar: 2μm. (F) Quantification of relative distance between Chr15 and Chr17 (left), and relative distance of Chr15 and Chr17 to the nucleus center (right) in WT and 25A haESCs. (mean ± SD, two-tailed ratio t-test, *: p<0.05, **: p< 0.01, ***: p< 0.005, ****: p< 0.0001, ns: not significant. n = 29 and 26 cells, respectively.)

In addition, based on the Hi-C results, we inferred consensus 3D structure of the genome in both WT and 25A cells (Figure 2C, Video S1, S2). Notably, Chr15 and Chr17, which were separated in WT, clustered together in 25A. Moreover, the territory of Chr17 was located close to the nucleus center in WT, but the corresponding moiety of Chr17 in the fusion Chr15-17 moved outward to the edge of the nucleus in 25A, similar to the radial location of the Chr15 moiety (Figure 2D), indicating that the radial position of the Chr17 relative to the center of the nucleus also changed significantly after chromosome fusion. Interestingly, most of the chromosomes stayed in their radial positions, but a few chromosomes showed radial displacements, such as Chr9 moved outward while Chr12 and 16 shifted inward (Figure 2D), which were likely a result of the disturbance caused by redistribution of Chr15-17 territory. Further 3D-FISH analysis in 25A cells consistently showed a juxtaposition of Chr15 (labelled with red) and Chr17 (labelled with green), an indication of a single territory of the fusion chromosome (Figure 2E). Statistical analysis also revealed that Chr15 and Chr17 clustered together in 25A and the radial position of Chr17 in the nucleus was shifted outward (Figure 2E, 2F). Notably, within the new single chromosome territory formed by fused Chr15-17, there was no extensive intermingling between the two moieties of Chr15 and Chr17 (Figure 2E), suggesting that a chromosome territory is not chaotically arranged, but rather likely determined by continuous DNA sequences and cis-interactions between chromatin loops or topologically associating domains within each chromosome.

Previous studies have suggested that larger chromosomes are more likely to be distributed at the periphery of the nucleus, while the shorter chromosomes tend to be located in the center (Bolzer et al., 2005; Boyle et al., 2001; Cremer et al., 2001; Mayer et al., 2005; Sun et al., 2000; Tanabe et al., 2002); the chromosome with higher and lower gene density are respectively located at the center and the edge of the nucleus (Boyle et al., 2001; Cremer et al., 2003; Cremer et al., 2001; Mayer et al., 2005). Consistent with previous reports, the fusion of Chr15 and Chr17, which are relatively short among the mouse native chromosomes, increased chromosome size (even longer than the largest Chr1) in 25A (Figure S2G-I), and the fusion chromosome is located at the edge of the nucleus (Figure 2D, Figure S2J). The radial position of a chromosome and gene density are negatively correlated in WT cells (p = 0.015). However, this correlation is disrupted in 25A (p = 0.158) (Figure S2J, S2K). The gene density of fused Chr15-17 was higher than the average gene density (Figure S2J, S2K), but it still moved outward, indicating a weak correlation between radial position of a chromosome and gene density in mouse cells.

### Chromosome territory rearrangement has little effect on gene expression

Chromosome territory as well as inter-chromosome interactions has been suggested to affect gene expression (Mahy et al., 2002; Osborne et al., 2004; Volpi et al., 2000; Williams et al., 2002). Thus, we performed RNA-seq and transcriptome analyses in 25A cells. To our surprise, although the fusion of Chr15 and Chr17 resulted in drastic changes in both the relative positions and the radial distributions of chromosome territories, these perturbations exerted no apparent effects on global gene expression. Compared to WT cells, only 0.33% of the genes in the whole genome of 25A cells displayed significant differential expression (FDR<0.05, log2(FC)>0.5) (Figure 3A, Figure S3A, S3B), most of which (94.8%) had an expression difference of less than 2-folds (Figure S3C). Interestingly, there was no correlation between the distribution of differentially expressed genes and the specific changes of chromosomal territories on Chr15 and 17 (Figure 3B, Figure S3D). These results indicated that the rearrangement of chromosome territories by chromosome fusion imposes little effect on gene expression.

**Figure 3.**
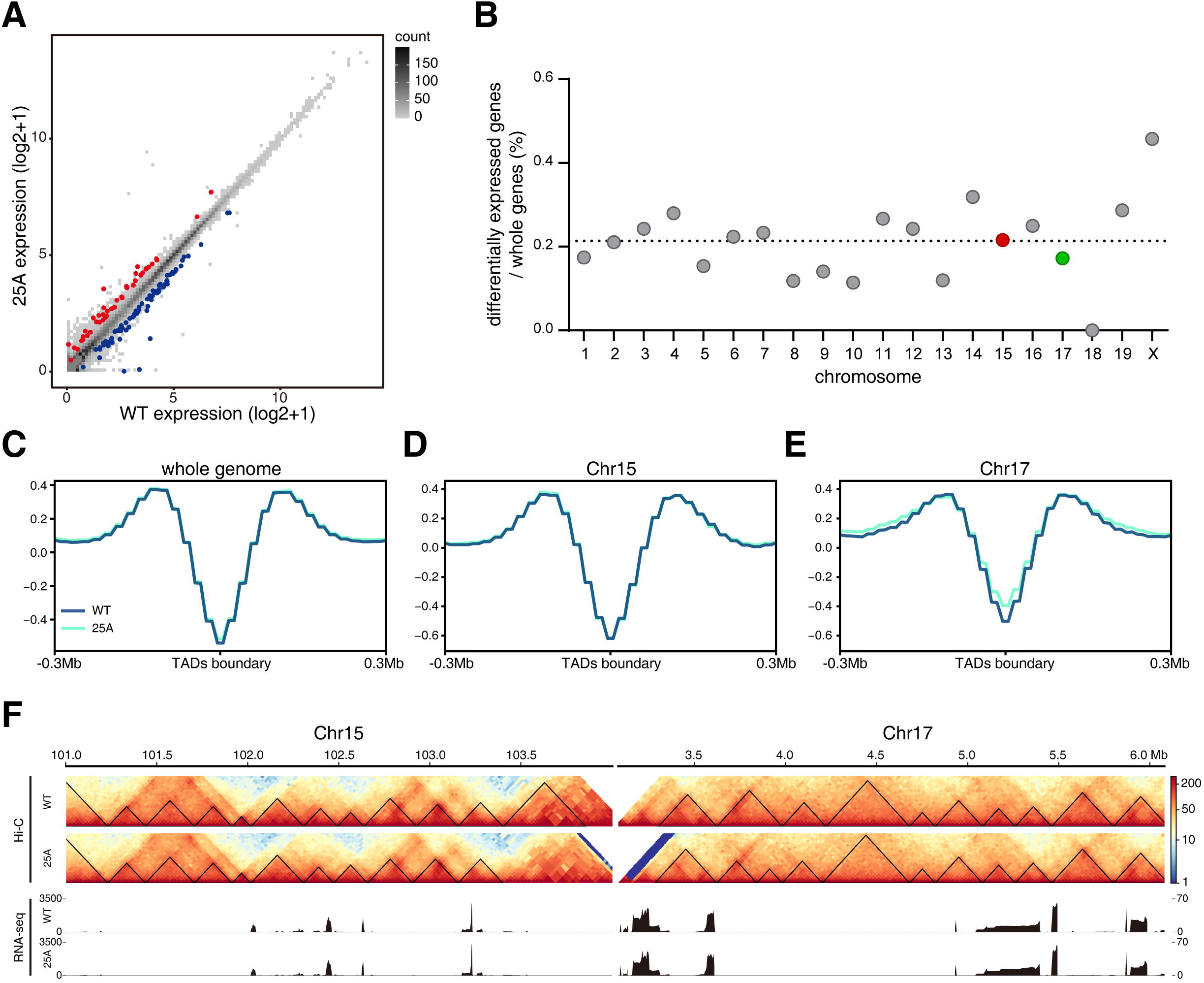
Transcriptome and TADs analyses in 25A cells. (A) Comparison of gene expression in 25A cells and WT cells. Up-regulated and down-regulated genes were shown in red and blue, respectively (FDR < 5%). (B) The proportion of the differentially expressed genes in each chromosome of 25A cells. The dash line indicates the average proportion of differentially expressed genes of the whole genome. (C to E) Comparison of TAD boundaries in WT and 25A cells. Aggregate profiles of insulation scores around TADs boundaries at a 10 Kb resolution for the whole genome (C), Chr15 (D) and Chr17 (E). (F) Hi-C contact matrixes of the 3 Mb region upstream from chromosome fusion site on Chr15 and downstream on Chr17 of WT and 25A cells. TADs are depicted as black triangles. RNA-seq expression tracks in the corresponding fusion regions in both WT and 25A cells are shown in lower panels.

The loose link between the significant reorganizations of chromosome territories and the subtle changes of gene expressions prompted us to ask whether chromosome fusion affected the lower levels of chromatin architecture, e.g. topologically associating domains (TADs), since TADs have been considered to be the functional units that regulate gene expression (Dixon et al., 2015; Dixon et al., 2012; Nora et al., 2012). Therefore, we further analyzed TADs in the whole genomes of both WT and 25A cells, and found that the insulation score near TADs boundaries across whole genome, including Chr15 and Chr17 in the WT cells and Chr15-17 in the 25A cells were indistinguishable (Figure 3C-E), indicating that the changes of chromosome territories did not disturb the overall TADs. Surprisingly, compared to that in WT, TADs within the 6-Mb fusion regions of Chr15 and Chr17 in 25A remained largely unchanged (Figure 3F, upper panels), providing a plausible explanation for nearly the identical gene expression patterns within the fusion regions in the WT and 25A cells (Figure 3F, lower panels).

### Generation of chromosome fusion mice

Chromosome territories appear to be different in different cell types (Mayer et al., 2005; Meaburn et al., 2016), suggesting that there are functional correlations between chromosome architecture (chromosome interaction) and gene expression. In order to explore further the effect of chromosome-fusion induced changes of chromosome territory at the organismal level, we audaciously injected 25A haESCs into WT mouse oocytes through intracytoplasmic AG-haESC injection (ICAHCI), and the resulting embryos were implanted into the uterus of surrogate mother mice (Figure 4A). Like the WT haESCs, 25A haESCs were functioning as “sperms”, and yielded heterozygous F0 female mouse, whose cells contained both the fusion Chr15-17 and the native Chr15 and 17 (Figure 5A). Karyotype analysis of the bone marrow cells confirmed that the F0 female mouse had a fused Chr15-17 and the mini-chromosome, which existed in 25A cell line (Figure S4B). The appearance and growth of F0 heterozygous female mice were not significantly different from the mice parallelly generated with WT haESCs through the ICAHCI method. We then examined the reproductive capability of F0 female mice by both *in vitro* fertilization (IVF) and natural cross-breeding with WT males, respectively, and both methods produced healthy F1 offspring with approximately 1:1 ratio of heterozygous to wild-type mice (Figure S4C), which fits the Mendelian genetics well. Interestingly, the mini-chromosome seen in the F0 mice was absent in heterozygous F1 mice (Figure 4C, Figure S4D). We speculated that the mini-chromosome might have been lost during oocyte meiosis in F0 mice. These results indicate that the Chr15-17 fusion is “overlooked” by the zygotes, and as a result, the reproduction of heterozygous female mice is smooth as usual.

**Figure 4.**
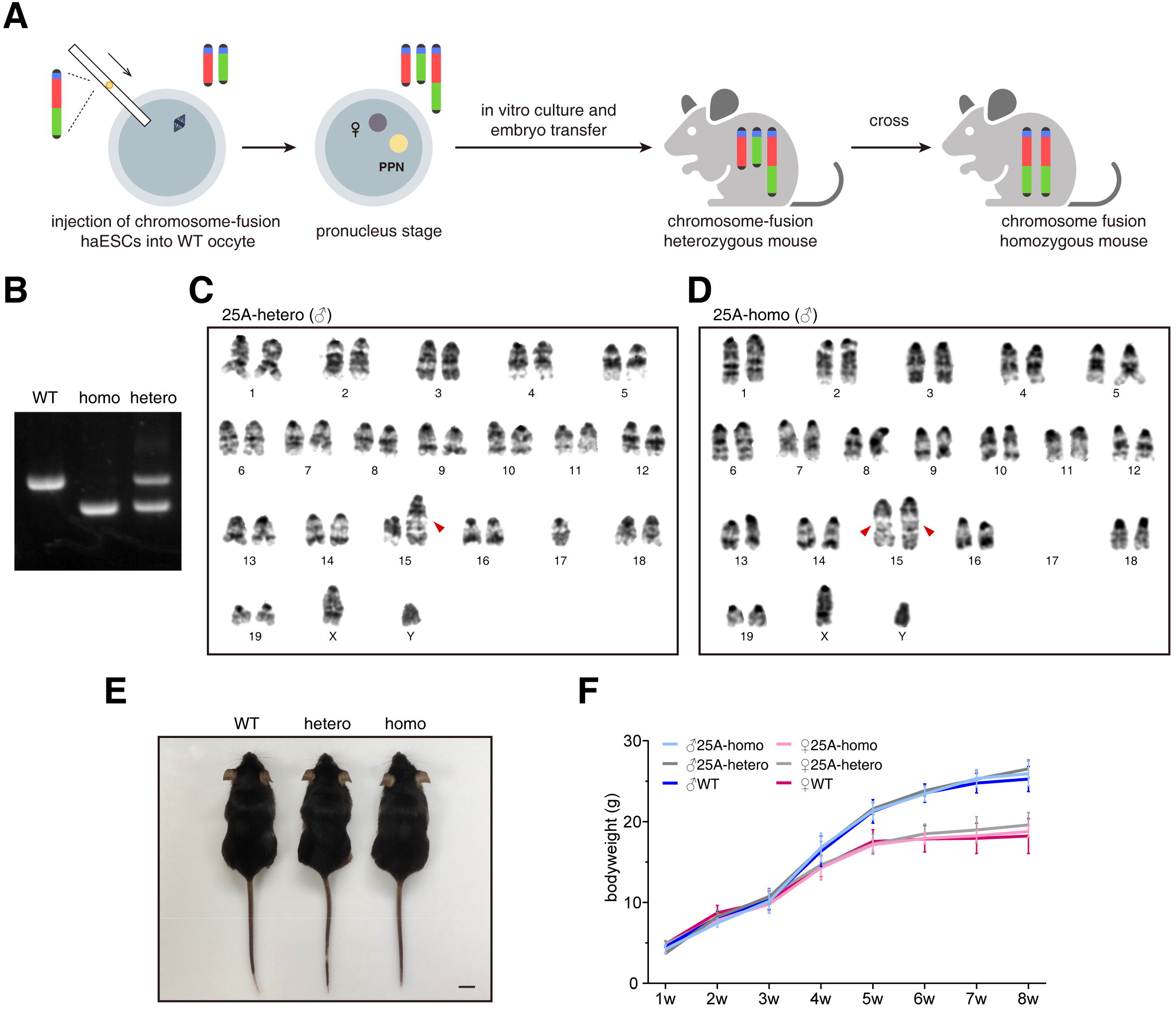
Generation of chromosome fusion mice. (A) Schematic procedures showing the generation of chromosome-fusion heterozygous mice (see Methods). Chromosome-fusion homozygous mice are generated by crossing between heterozygous female and male mice. PPN, pseudo-pronucleus derived from the injected haESCs. (B) Genotype analysis of the chromosome fusion homozygous and heterozygous mice. WT and homozygous mice showed a ~860 bp and a ~660 bp band, respectively, while heterozygous mouse showed both. (C) G-band karyotype analysis of 25A heterozygous male mouse (37+XY, t(15;17)(F3;A2)). Red arrowhead indicates the fused Chr15-17. (D) G-band karyotype analysis of 25A homozygous male mouse (36+XY, t(15;17)(F3;A2)×2). Red arrowhead indicates the fused Chr15-17. (E) The appearances of a WT male mouse (left), a 25A heterozygous male mouse (middle) and a 25A homozygous male mouse (right). (F) Growth curve along postnatal development of the WT, 25A heterozygous and homozygous mice from 1 week to 8 weeks.

**Figure 5.**
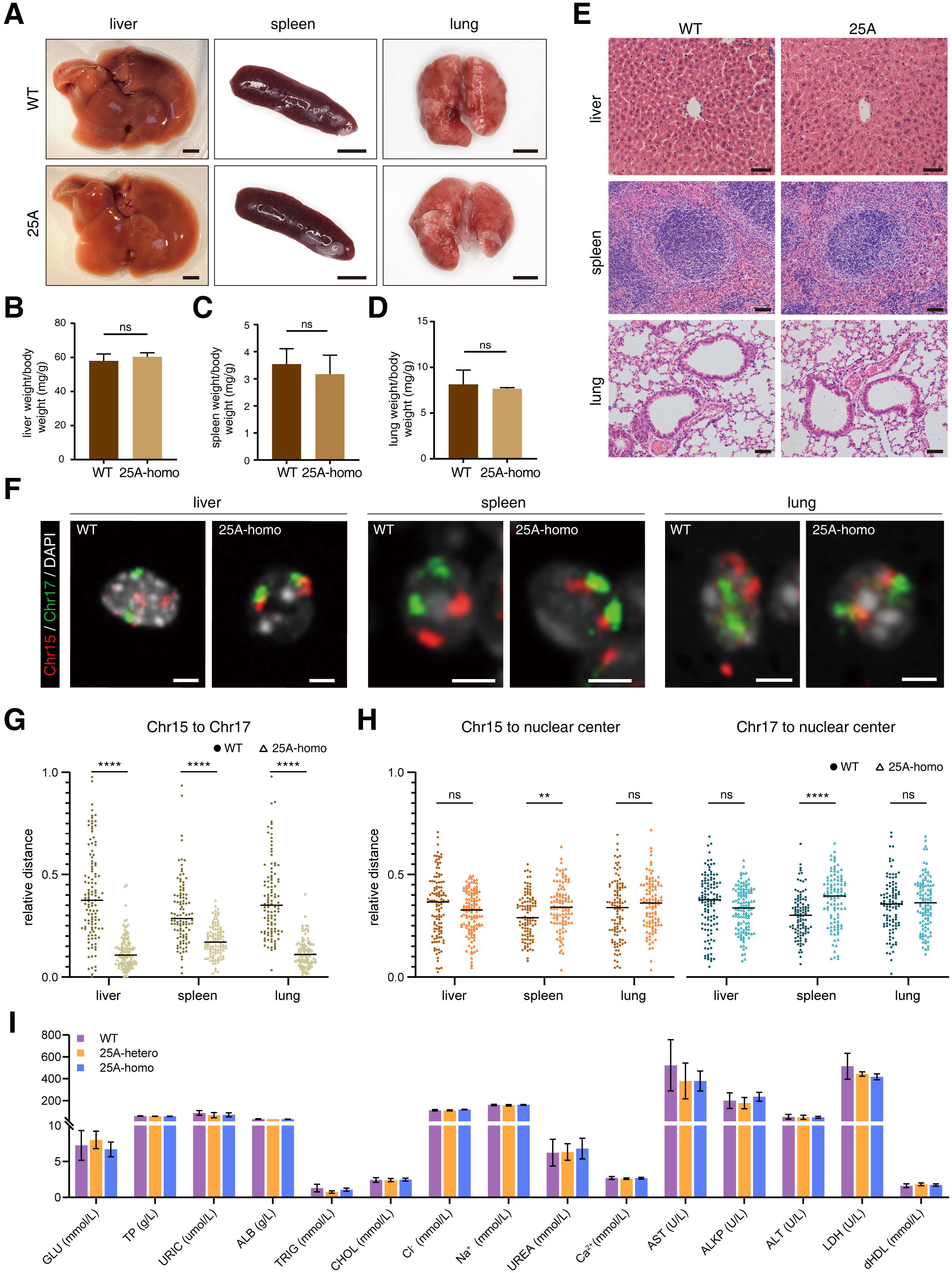
Phenotypic analyses of chromosome fusion mice. (A) Morphological features of liver, spleen and lung from adult WT and 25A (Chr15-17 fusion) homozygous mice. Scale bar: 300 mm. (B to D) Relative organ weight to body weight of liver (B), spleen (C) and lung (D) in 8 weeks WT and Chr15-17 fusion homozygous mice. (mean±SD, two-tailed t-test, n=3). (E) H&E staining of liver, spleen and lung from 8 weeks WT and Chr15-17 fusion homozygous mice. Scale bar: 50 μm. (F) Chromosome FISH for Chr15 (red) and Chr17 (green) in the cells of liver (left), spleen (middle) and lung (right) from WT and Chr15-17 fusion homozygous mice. The nuclei were stained with DAPI (grey). Scale bar: 2.5 μm. (G) Quantification of relative distance between Chr15 and Chr17 in the cells of the corresponding organs of WT and Chr15-17 fusion homozygous mic e. (Two-tailed t-test, n=118, 120, 102, 102, 104 and 104 chromosomes, respectively.) (H) Quantification of relative distance of Chr15 and Chr17 to the nucleus center in the cells of the corresponding organs of WT and Chr15-17 fusion homozygous mice. (Two-tailed t-test, n=118, 120, 102, 102, 104 and 104 chromosomes, respectively.) (I) Serum biochemical test of WT, Chr15-17 fusion heterozygous and homozygous mice. (mean±SD, n=5, 3, 3 mice, respectively.)

We took a step further to cross heterozygous mice and obtained homozygous mice containing two copies of the fusion chromosomes (Figure 4A, 4B). Karyotype analysis of the homozygous mice showed two fused Chr15-17 (Figure 4D, Figure S4E). Apparently, homozygous, heterozygous and WT mice showed no differences in appearance, postnatal growth and development (Figure 4E, 4F). Homozygous male and female mice can produce homozygous offspring, and their reproductivity is nearly the same as that of WT mice. These results indicate that chromosome fusion has no obvious effect on the growth, development and reproduction of mouse.

### Chromosome territory rearrangement by chromosome fusion causes no detectable defects in mouse tissues or organs

We then examined the effects of chromosome fusion on the functions of different tissues and organs in the homozygous and heterozygous mice. The shape and weight of the main organs (such as liver, spleen and lung) in the heterozygous and homozygous mice were very similar to those of WT mice (Figure 5A-D, Figure S5A, S5B). Additional HE staining showed that both the internal structure and the constitutions of the main organs (liver, spleen, lung, heart and kidney) as well as reproductive organs (testis and ovary) matched well with those of WT mice (Figure 5E, Figure S5C-E). Further chromosome FISH on the cells of several organs revealed that in homozygous mice, the Chr15-17 fusion was intact, and the rearranged chromosome territories were different from those observed in the cells from WT organs (Figure 5F, 5F). Notably, the radial position of Chr15-17 showed outward shift in spleen cells, but no significant movement in liver and lung cells (Figure 5F, 5H), consistent with the notion that a given chromosome in different cell types may display different territories (Cremer et al., 2001; Croft et al., 1999; Meaburn and Misteli, 2007). Regardless of cell-type differences, within the single territory of fused Chr15-17, the moieties of Chr15 and Chr17 distal to the fusion regions did not mingle with each other (Figure 5F), further supporting the model that a chromosome territory is primarily determined by continuous DNA sequences and the cis-interaction between chromatin loops or TADs with special proximity. At the metabolic level, the blood parameters, including complete blood count and serum biochemical test, were analyzed, and found no significant differences between WT and chromosome-fusion mice (Figure 5I, Table S1). We thus concluded that the rearrangements of chromosome territories by chromosome fusion do not cause detectable defects in mouse cell differentiation, organogenesis and development.

## Discussion

There are twenty chromosomes in the haploid of mouse cells, nineteen autosomes (from Chr1 to Chr19) and one sex chromosome (ChrX or ChrY). The autosomes are likely numbered according to their size: Chr1 is the longest (195 Mb), and Chr19 is the shortest (62 Mb). The lengths of X and Y chromosomes are 169 Mb and 91 Mb, respectively (Figure S2I). Mouse autosomes are all telocentric, i.e. the centromere of the chromosome is located next to one of the telomeres (Figure S1A). Though the DNA in both centromeric and telomeric regions are repetitive sequences (Garagna et al., 2002; Kalitsis et al., 2006), the primary structure of a centromere consists of minor satellite and major satellite, and is much more complicated than that of a telomere, which mainly consist of regular (TTAGGG)_n_ repeats (Figure S1A) (Kalitsis et al., 2006). The telomere-centromere layout of mouse chromosome facilitates chromosome engineering: deletions of both centromere and the proximal telomere in a given chromosome can be done at one CRISPR-Cas9 cut (Figure 1A, Figure S1A). However, poor annotations of the genes near each centromeric region lead to potential uncertainties of chromosome fusion. We first carefully analyzed all of the genes near centromere in every chromosome, and found that some of the chromosomes, such as Chr 13, 15, 16 and 17 are suitable candidates for chromosome fusion.

Second, chromosome size after fusion could be potentially problematic, because there might be a length limit that a cell can tolerate (Hudakova et al., 2002; Rens et al., 2006; Schubert and Oud, 1997). Third, the radial distribution of chromosome territories in the nucleus have been proposed to be correlated with chromosome length and gene density (Boyle et al., 2001; Cremer et al., 2003; Cremer et al., 2001; Mayer et al., 2005). Chromosomes with larger size and lower gene density tend to be distributed at the periphery of the nucleus, and *vice versa*. Given that (1) Chr15 and Chr17 are relatively small, and their fusion results in a 199-Mb chromosome, which is slightly longer than the largest Chr1 (195-Mb) (Figure S2I); (2) the gene density of Chr17 is significantly higher, while the gene density of Chr15 is lower than that of other chromosomes, and the fused Chr15-17 displays a medium gene density between the Chr15 and Chr17 (Figure S2I), we have chosen Chr15 and 17 to perform chromosome fusion. In spite of these concerns, the successfully construction of fusion chromosome implies that reconstruction of mouse genome might not be an impossible task.

The eukaryotic genome seems to have evolved into a hierarchical structure, including chromatin loops, TADs, compartments and chromosome territories (Cremer and Cremer, 2001; Lamond and Earnshaw, 1998; Szczepinska et al., 2019). The driving forces for these hierarchical arrangements are still mysterious. It has been proposed that chromosome territory regulates genome architecture and thereby affects gene expression (Mahy et al., 2002; Osborne et al., 2004; Volpi et al., 2000; Williams et al., 2002). However, in the single cell organisms like *S. cerevisiae* and *S. pombe*, drastic chromosome architecture changes by artificial chromosome engineering cause marginal changes of gene expressions, and affect little the functions of yeast cells (Gu et al., 2022; Luo et al., 2018; Shao et al., 2018). In addition, the chromosome fusion in haESCs results in chromosome distribution changes in cell nucleus, but neither induces obvious genome instability, nor causes detectable defects in gene expression, pluripotency and gamete function (Figure 1 to Figure 3). Importantly, the mice containing the fusion chromosome are apparently healthy, and competent to propagate regardless of the chromosome territory perturbations in various cell types of individual tissues and organs (Figure 4 and Figure 5). These lines of evidence strongly suggest that the territories of individual chromosomes in eukaryotic nucleus might be passively demarcated under the scenarios of geometric constraints and chromatin collisions in the limited volume of cell nucleus, and the effects of chromosome territories on gene expression are generally insignificant or even dispensable. However, there are cases that trans-interactions between two chromosomes are of functional (Branco and Pombo, 2006b; Maass et al., 2019; Spilianakis et al., 2005), suggesting coincidences of evolution. Nevertheless, the chromosome fusion does not seem to change TADs of the corresponding chromosomes (Figure 3C-E), consistently supporting the hypothesis that TADs are the functional units for the regulation of gene expression (Dixon et al., 2015; Dixon et al., 2012; Nora et al., 2012).

Different species on earth have different numbers of chromosomes. It remains elusive whether the genome organization in different species is randomly or fortuitously retained during long course of evolution. We have arbitrarily fused Chr15 and 17 in haESCs, and fortunately cultivated the mice with 2n=38, indicating the high plasticity of mouse genome. The Chr15-17 fusion disturbs the radial positions of territories of the natural Chr15 and 17 (Figure 3C-F), as well as Chr9, 12 and 16, but not others (Figure 3D). Why and how the regional territory perturbations only affect some of the chromosomes remains unclear. In addition, we do not know whether the haESCs used in this work are able to tolerate additional chromosome fusions, and still able to maintain its pluripotency and gamete functions afterwards. Ideally, more dramatic changes of chromosome territories rely on more massive chromosome engineering, for example, to construct n=18 (or even less) haESCs and/or mice through three-chromosome fusion or two-pairs of chromosome fusion, or fusion between long chromosomes. Further extensive karyotype engineering will help to further clarify the structure-function correlations between chromosome territories and genome activities. It seems to be technically feasible, but the global orchestration of genomic activities and regulations within the functional milieu of the mammalian cell nucleus is so sophisticated that further extensive karyotype engineering might be extremely challenging.

## Supporting information

Video S1. 3D genome reconstuction of WT

Video S2. 3D genome reconstuction of 25A

## Acknowledgments

This research was supported by grants from National Key Research and Development Program of China (2019YFA0109902 to J.-Q.Z) and Chinese Acedamy of Sciences (ZDBS-LY-SM018 to J.-Q.Z. and Y.Z.).

## Author contributions

J.-Q.Z. and Y.Z. conceived and supervised the project, analyzed the data, and drafted the manuscript. Y.W. constructed the chromosome fusion in haESCs, carried out most of the experiments and data analyses, and drafted the manuscript. Z.Q. and Y.F. conducted IHC staining and some of the histological experiments and drafted the manuscript. Y.C. and J.S. performed Hi-C and RNA-seq data analyses. J.P. and J.S. provided help with the histological experiments. J.L. provided the original haESCs and contributed to data analyses.

## Supplementary Information

Supplementary Information is available for this paper.

## Materials and Correspondence

Correspondence and requests for materials should be addressed to Y.Z.

## Materials and Methods

### Animal Use and Care

All specific pathogen-free (SPF)-grade mice were maintained and handled in accordance with the ethical guidelines of the Center for Excellence in Molecular Cell Science, Chinese Academy of Sciences (CAS). The uses of experimental animals were maintained and handled in strict accordance C57BL/6 mice used for mating and propagation, and BABL/c nude mice used for teratoma formation were obtained from Shanghai Jihui Laboratory Animal Care Company.

### Cell culture

haESCs (H19ΔDMR-IGΔDMR-AGH) were maintained in a standard ESC culture system: DMEM (Millipore) with 15% FBS (Gibco), penicillin-streptomycin (Gibco), nucleosides (Millipore), non-essential amino acids (Millipore), L-glutamine (Millipore), β-mercaptoethanol (Millipore), 1,000□JU/ml Lif (Millipore), 3□JμM CHIR99021 (Selleck) and 1□JμM PD03259010 (Selleck) (Qu et al., 2018; Yang et al., 2012; Ying et al., 2008; Zhong et al., 2015).

### Fluorescence-activated cell sorting (FACS)

haESCs were trypsinized into single cells and incubated with 15□Jμg/ml Hoechst 33342 (Invitrogen) in a 37 °C water bath for 5 min. Sorting was then conducted to harvest the haploid 1n peak by using FACS Aria II (BD Biosciences) (Leeb and Wutz, 2011; Qu et al., 2018; Zhong et al., 2016; Zhong et al., 2015).

### CRISPR-Cas9 fused chromosomes in haESCs

The sgRNAs of chromosome 15 and chromosome 17 were connected to the pX330-mCherry plasmid (Addgene, 98750). WT cells were transfected with 250□Jµl Opti-MEM that contained 5□Jµl Lipofectamine 2000 (Thermo Fisher) and□J2.5µg sgRNA-pX330-mCherry plasmid. 20–48 h after transfection, haploid cells expressing red fluorescent protein were enriched by FACS and plated into a well of a 6-well plate at a low density of around 4000 cells per well. Single colonies were picked and passaged to a well of a 96-well plate after 5-8 days. Related sequences are listed in Supplementary Table 2.

### Cell proliferation

Haploid cells enriched by FACS were collected to evaluate cell proliferation rate, 4.5 × 104 sorted cells were cultured in a well of a 24-well plate. After 3 days, cells were dissociated and counted.

### Intracytoplasmic AG-haESCs injection (ICAHCI) and embryo transfer

ICAHCI and embryo transfer to generate mice were performed with the help of the Animal Core Facility, the Center for Excellence in Molecular Cell Science, Chinese Academy of Sciences (CAS) as described previously (Yang et al., 2012; Zhong et al., 2015).

### Karyotype Analysis and cell FISH

haESCs were incubated with 0.4 mg/ml demecolcine (Sigma) for 1 h. After trypsinization, the cells were resuspended in 0.075 M KCl at 37°C for 15min and then fixed in methanol: acetic acid (3:1 in volume) for 30min. The cells were dropped onto pre-cold and precleaned slides. For karyotype analysis, the protease-treated cells were stained with Giemsa dye (Yeasen) for 15min. Pictures were taken by Olympus BX53 and more than 50 metaphase spreads were analyzed. The G-banded ideogram of chromosome images was arranged according to the previous publication (Mann et al., 1990).

For cell FISH experiments, whole chromosome probes XMP 15 and XMP17 were hybridized following the manufacturer’s protocol (MetaSystems), and nuclei were counterstained with DAPI. Pictures of the chromosomes were acquired by using Leica TCS SP8 WLL. Only the stained chromosomes were analyzed.

### 3D FISH and image analysis

Cells grown on glass slide for 2 h were fixed with 4% PFA for 15min, permeabilized with 0.5% Triton in PBS for 20min and then in 0.1M HCl for 5min. The cells were then washed with 2×SSC for 5min twice and then washed in 50% formamide/4×SSC for 10h at 4°C. The hybridization was the same as mentioned above.

The images were acquired on Leica TCS SP8 WLL. For each imaging view, z-stacks covering the whole nuclei with a step size of 400 nm were taken for each channel and imaging conditions were kept for different views of one sample. DAPI was stained to represent the nuclear profile. The three-dimensional image analysis was carried out in Imaris (Bitplane) by ImarisCell, a module designed specifically to identify, segment, track, measure and analyze cell, nucleus and vesicles in 3D images. For 3D chromosome FISH image analysis, “Surface” function was used to segment nuclear boundary by DAPI channel and chromosome territory boundary of chromosome 15 and chromosome 17 by 488 nm and 552 nm channel intensity. The volume and center of mass of nucleus and chromosome territories were output directly. The volume of each nucleus was measured to normalize the volume of chromosome territories. Distance between nuclear center of mass and chromosome territories was normalized by the cubic root of nuclear volume. We only selected haploid cells for chromosome FISH analyses.

### Tissue FISH

The mice were euthanized by CO2 overdose. The tissues were harvested and fixed in 4% paraformaldehyde (PFA) and embedded in paraffin. After dewaxing and rehydration, the tissue section slides were heated in ddH2O for 25min and digested with pepsin (1mg/mL in 10mM HCl), and then washed in 50% formamide/4×SSC for 10h at 4°C. The XMP 15 and XMP 17 probes were added for hybridization at 80°C for 4min on a hot plate and then 37°C overnight in a humidified chamber. The glass coverslips were removed and the slides were washed in 0.1% tween-20/2×SSC at 37°C for 5min, and then 0.3% tween-20/0.4×SSC at 73°C for 2.5min. After draining, the slides were then washed in 0.1% tween-20/2×SSC at room temperature (RT) for 1.5min. Subsequently, the slides were briefly rinsed in ddH2O and then air dried at RT. Finally, nuclei were counterstained with DAPI. Pictures of the chromosomes were acquired by using Leica TCS SP8 WLL.

Quantification of relative distance of chromosome territories to the nuclear center as well as the relative distance between chromosome territories was done with ImageJ software. The center and area of each nucleus and chromosome territory of chromosome 15 and 17 were measured. Relative distance between chromosome 15 and 17 was calculated as the shortest distance between two pairs of chromosome 15 and 17 in diploid cells and was normalized by the square root of nuclear area.

### Hematoxylin & Eosin staining

Animals were sacrificed and tissues were harvested and fixed in 4% PFA, embedded in paraffin. After dewaxing and rehydration, tissue section slides were stained with hematoxylin stain solution (Yeasen) for 5min and eosin Y stain solution (Yanye) for 10s. The slides were dehydrated in increasing concentrations of alcohols, cleared by xylene, and mounted in neutral balsam. Pictures were taken by Olympus BX53.

### Teratoma formation and immunohistochemistry

The Diploid (2n) cells of haESCs were purified with FACS. Di-haESCs (approximately 1 × 107 cells) were trypsinized into single cells with PBS buffer and subcutaneously injected into BABL/c nude mice (4 weeks old). Ten mice were injected for each cell line. After 4-6 weeks animals were sacrificed and teratomas were fixed in 4% PFA, embedded in paraffin.

For immunohistochemistry, teratoma sections were blocked using 10% normal goat serum (Solarbio) in PBS for 1h and stained with primary antibody at 4°C overnight, followed by secondary antibody (SCBT) for 1h at RT and DAB enhancer (MKbio). After hematoxylin staining for 2min, the slides were dehydrated in increasing concentration of alcohols, cleared by xylene, and mounted in neutral balsam. Pictures were taken by Olympus BX53.

The used primary antibody: anti-alpha-1-fetoprotein (ARG56134, Arigobio) at 1:500 dilution for endoderm lineage; anti-smooth muscle actin (sc-53142, SCBT) at 1:50 dilution for mesoderm lineage; anti-Tuj1 (sc-80005, SCBT) at 1:400 dilution for ectoderm.

### Measurements of blood parameters

The blood samples were collected into micro blood collection tubes by the retro-orbital bleeding in mice.

Microtubes contained EDTA as anticoagulants to be used for hematological examinations, and microtubes without EDTA were used for clinical chemistry serum measurements. All samples received hematological parameters on XN-1000V Hematology Analyzer (Sysmex). The clinical chemistry parameters glucose (GLU), total protein (TP), uric acid (URIC), albumin (ALB), triglyceride (TRIG), cholesterol (CHOL), chloride (Cl-), sodium (Na+), urea, calcium (Ca), aspartate aminotransferase (AST), alkaline phosphatase (ALKP), alanine aminotransferase (ALT), lactate dehydrogenase (LDH), and high-density lipoproteins (HDL) were determined in the serum with a VITROS 4600 (Ortho Clinical Diagnostics).

### Hi-C library preparation and sequencing

Cells were cross-linked with 3% fresh formaldehyde (final concentration) and then quenched with 0.15 M glycine (final concentration) for 5min. Genomic DNA was extracted and digested with 200 units MboI (NEB) (Lieberman-Aiden et al., 2009). DNA ends were labeled with biotin-14-dCTP (TriLINK), and after ligated, they were sheared to a length of ~400 bp. Point ligation junctions were pulled down with Dynabeads MyOne Streptavidin C1 (Thermo). The Hi-C library for Illumina sequencing was prepared with the NEBNext Ultra II DNA Library Prep Kit for Illumina (NEB) according to the manufacturer’s instructions. Paired-end sequencing (150 bp read length) was performed on the NovaSeq 6000 platform (Illumina) and 400 Gb raw reads were obtained. Paired-end sequencing reads were trimmed for adaptors and low-quality reads by fastp (v0.21.0) with default parameters (Chen et al., 2018). Trimmed reads were processed using HiCExplorer (v3.7.2) as previously described (Ramirez et al., 2018). Briefly, mates were mapped individually to Mouse genome GRCm39 using BWA (bwa-0.7.17) with the option ‘mem -A 1 -B 4 -E 50 -L 0’, the resulting BAM files were used to build contact matrices at binning resolutions of 20 kb and 1 Mb. The raw contact matrices were normalized using a fast balancing algorithm introduced by Knight and Ruiz (KR) (Knight and Ruiz, 2013) to correct bias and scaled to a fixed read count defined by the sample with the lowest coverage. TADs were identified using command ‘hicFindTADs’ with default parameters, the resulting bedGraph files with insulation scores were processed by deepTools (v3.4.3) for further visualization (Ramirez et al., 2016).

### Inferring the 3D structure of the genome

The 3D structure of the genome in WT and 25A cells were inferred using PASTIS v0.5 with the PM2 algorithm (Varoquaux et al., 2014), the resulting Protein Data Bank (PDB) files were visualized with Pymol (v2.4.0).

### RNA-seq analysis

Total RNA was isolated from the cells using Trizol reagent (Ambion). The library preparation followed the standard procedure (Illumina). The libraries were sequenced on the Illumina NovaSeq 6000 platform using the 150-bp paired-end sequencing strategy. For each sample, 8 Gb clean data was obtained. The clean reads were mapped to the reference genome GRCm39 using STAR (v2.0), the resulting BAM files were converted to genome-wide tracks using deepTools. Transcript quantification was performed using kallisto (v0.46.1) with default parameters, which was further processed by DESeq2 (v1.14.1) for differential expression with a FDR cutoff of 5%.

### EB formation and differentiation of haESCs

Before EB formation, the dishes were coated with gelatin. The haESCs were trypsinized and diluted into 2.5 × 105 cells/ml in the differentiation medium (DM: DMEM with 20% FBS, non-essential amino acids, β-mercaptoethanol, L-glutamine, penicillin/streptomycin, and sodium pyruvate), and aliquoted into 20 μl drops on the lid of 10 cm culture dishes, and 50 drops in total following the standard hanging-drop method (Boheler et al., 2002; Wobus et al., 2002). The droplets were collected from the lid on the next day and placed in 10 cm culture dishes filled with 10 mL DM and incubated at 37°C. The EBs were harvested 5 days later and transferred onto a new 48-well (gelatin coated) at a density of 10 EBs per well in DM and refresh the DM every 2 days. The differentiated EBs were harvested on the 14th day.

### Real-time PCR

Total RNA was extracted from the haESCs using TRIzol Reagent (Ambion). cDNA was generated by reverse transcription using Hiscript III RT supermix (Vazyme) according to the manufacturer’s instructions. The pluripotency marker genes were quantified by real-time PCR using AceQ Universal SYBR qPCR Master mix (Vazyme) and Roche LightCycler 96 qPCR Real-Time PCR system. Primers specific to each marker gene were listed in Supplementary Table 3.

## Supplemental information

Figure S1 (related to Figure 1). Genome stability and pluripotency of the chromosome fusion haESCs.

Figure S2 (related to Figure 2). Chromosomal interactions and territories are changed in 25A cells.

Figure S3 (related to Figure 3). Transcriptome analyses of 25A cells.

Figure S4 (related to Figure 4). Validation of heterozygous and homozygous mice containing fusion Chr15-17.

Figure S5 (related to Figure 5). Phenotypic analyses of 25A mice.

Table S1 (related to Figure 5). Complete blood counts of 25A mice.

Table S2 (related to Figure 1 and Figure S1). CRISPR-Cas9 target sites.

Table S3 (related to protocol). List of primers for PCR and real-time PCR.

Video S1 (related to Figure 2). 3D genome reconstuction of WT.

Video S2 (related to Figure 2). 3D genome reconstuction of 25A.

