## supplementary files for "Chromosome territory reorganization through artificial chromosome fusion is negligible to cell fate determination and mouse development"

### Figure S1

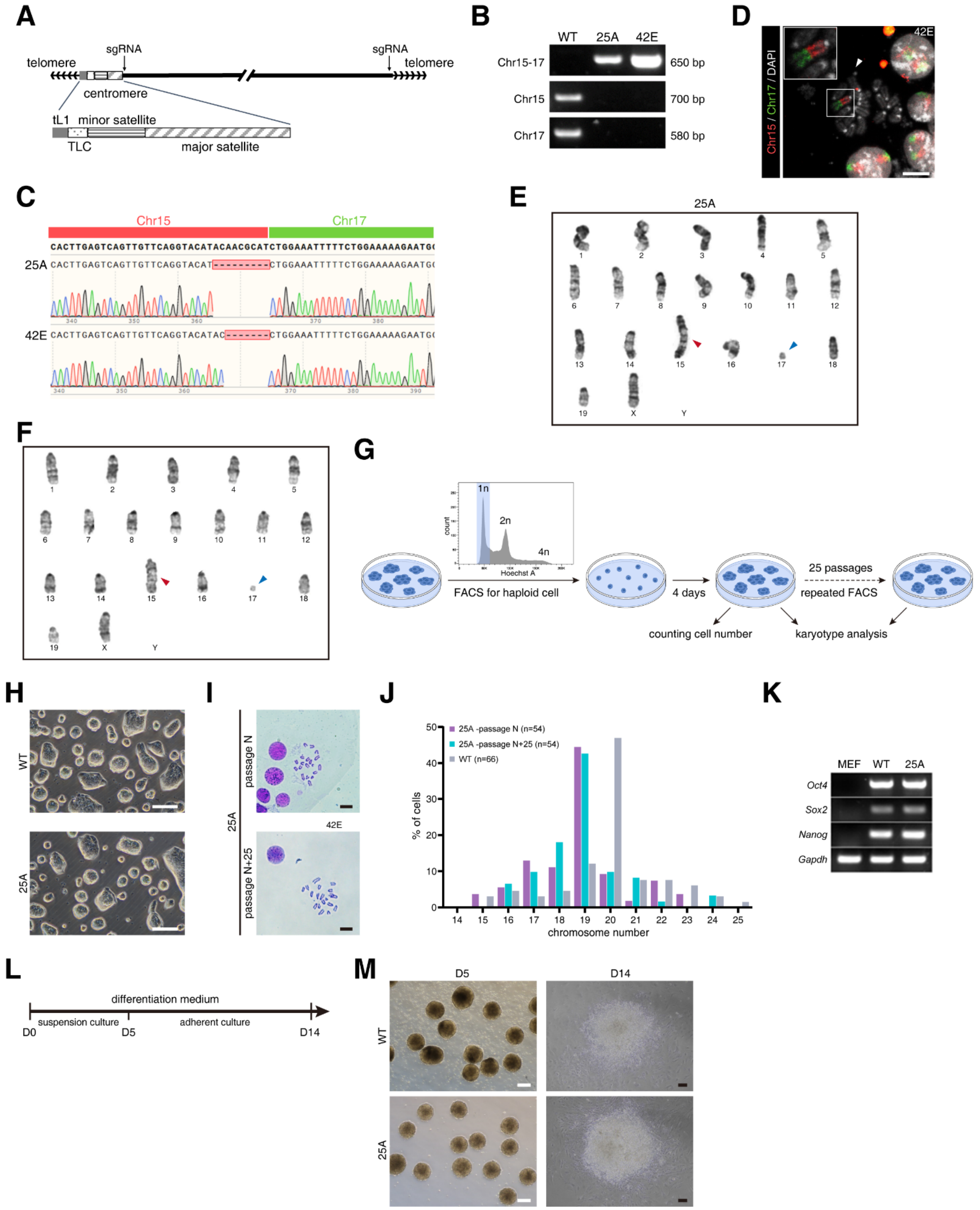

**Figure S1 (related to Figure 1). Genome stability and pluripotency of the chromosome fusion haESCs.**

(A) The typical structure of mouse telocentric chromosomes showing the positions of two telomeres, truncated LINE-1 (tL1), telocentric tandem repeat (TLC), centromeric minor satellite, and pericentric major satellite DNAs. The sgRNA target sites are near D-telomere region of chromosome 15 and C-telomere region of chromosome 17.

(B) PCR analysis of 25A and 42E haESCs. Cross-chromosomal PCR analysis with primer pairs 'F1' on Chr15 and 'R2' on Chr17, showed a ~650 bp band in 25A and 42E, while inter-chromosomal PCR detecting with 'F1' and 'R1' on Chr15 (~700 bp), and 'F2' and 'R2' on Chr17 (~580 bp) showed the expected bands only in WT.

(C) The sequencing results of the PCR products from the cross-chromosomal PCR in 25A and 42E haESCs, in which nine and seven bases were deleted at the junction site, respectively.

(D) Fluorescent images of the metaphase chromosomes of 42E haESCs labelled with Chr15 (red) and Chr17 (green) whole painting probes. Insets zoomed-in showing the fused Chr15-17 in 42E. The mini-chromosome was indicated with white arrowhead. Scale bar: 10 μm.

(E and F) G-band karyotype analysis of 25A (E) and 42E (F) haESCs: 19+X, t(15;17)(F3;A2) (red arrowhead), residual mini-chromosome (blue arrowhead).

(G) Diagram for assessing cell proliferation rate and karyotype stability. Haploid 25A cells are enriched by FACS (1n), and cultured in 2i medium for 4 days to count cell number. Karyotype analysis is performed before and after long-term culturing (25 passages).

(H) The colony morphology of WT (up) and 25A (down). Scale bar: 200 μm.

(I) Karyotype analysis of 25A haESCs of passage-N (up) and passages N+25 (down) showing n=19 and a mini-chromosome. Scale bar: 10 μm.

(J) Statistic analysis of chromosome numbers of WT (grey histogram) and 25A haESCs that passaged for N generations (purple histogram) and N+25 generations (blue histogram). The mini-chromosome was not counted in 25A.

(K) RT-PCR analysis of the pluripotency marker genes (*Oct4*, *Sox2* and *Nanog*) in WT and 25A haESCs. MEF cells and *Grapdh* gene were used as controls.

(L) Schematic of *in vitro* differentiation of haESCs to three germ layers.

(M) Differentiation morphologies of WT (up) and 25A (down) cultured in differentiation medium. Suspended embryoid bodies at day 5 (left) and attached morphology at day 14 (right) were shown. White scale bar: 500 μm; black scale bar: 100 μm.

**Figure S2**

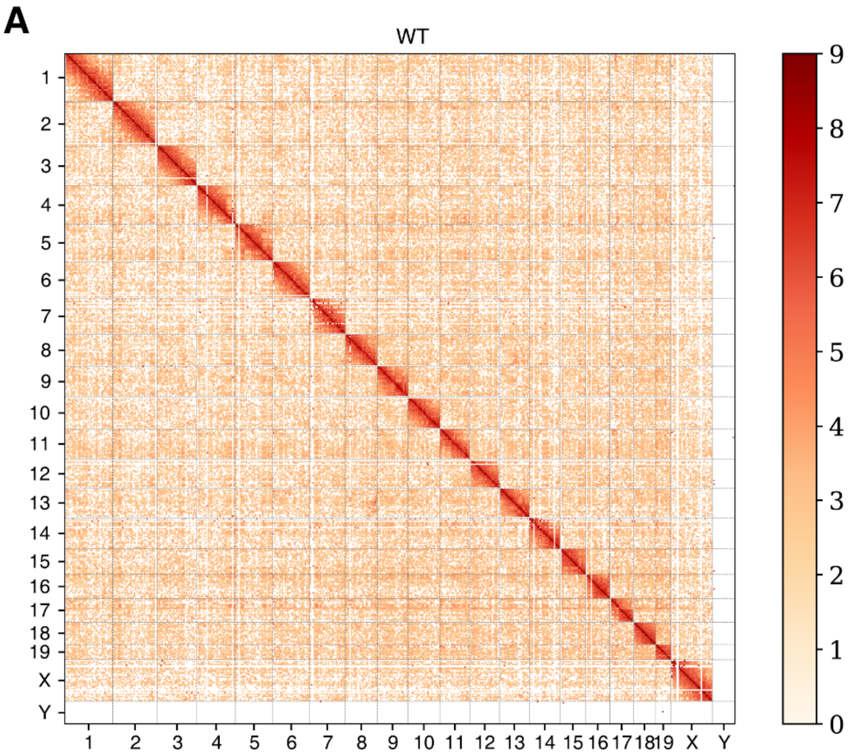
A
WT
X
Y
0
1
2
3
4
5
6
7
8
9

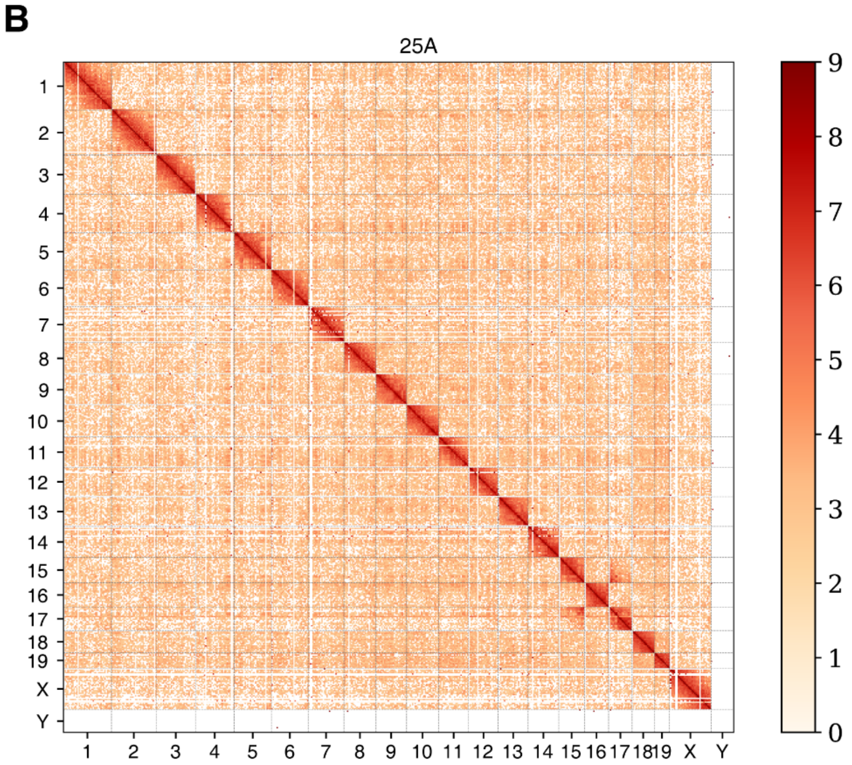
B
25A
X
Y
0
1
2
3
4
5
6
7
8
9

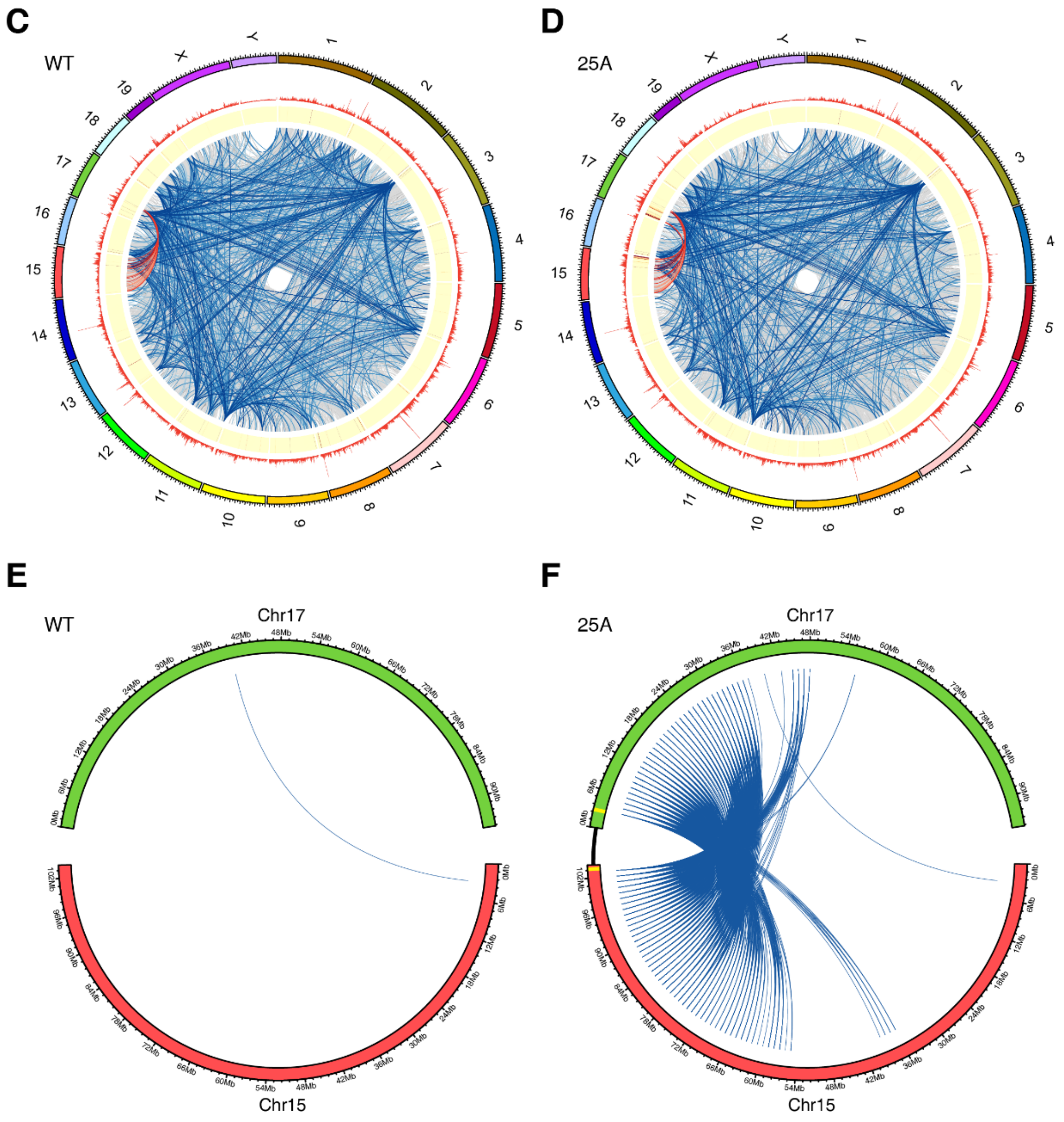
C
WT
D
25A
X
Y
E
WT
F
25A
Chr17
Chr15

G

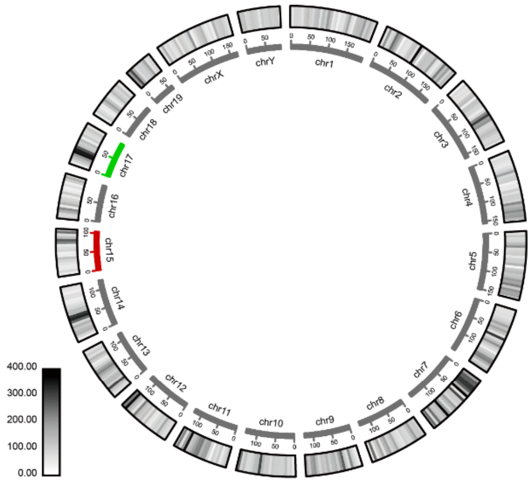
chr1
chr2
chr3
chr4
chr5
chr6
chr7
chr8
chr9
chr10
chr11
chr12
chr13
chr14
chr15
chr16
chr17
chr18
chr19
chrX
chrY
400.00
300.00
200.00
100.00
0.00

H

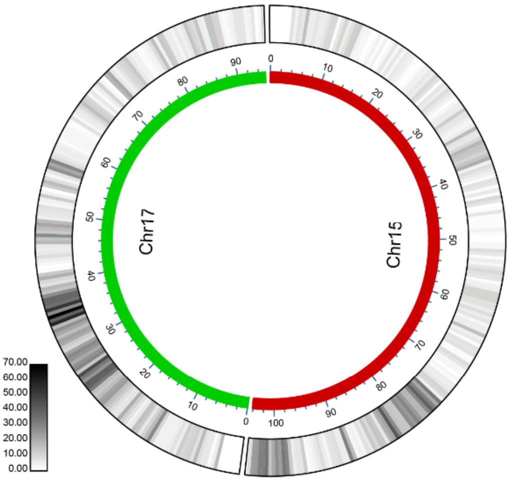
Chr17
Chr15
70.00
60.00
50.00
40.00
30.00
20.00
10.00
0.00

I

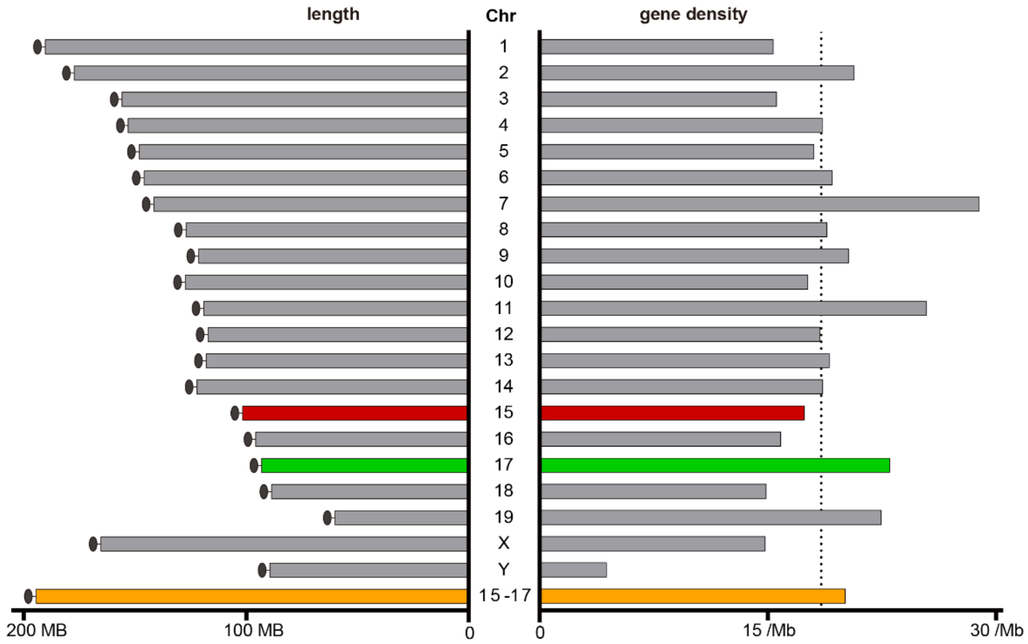
length
Chr
gene density
X
Y
15-17
200 MB
100 MB
0
0
15 /Mb
30 /Mb

J

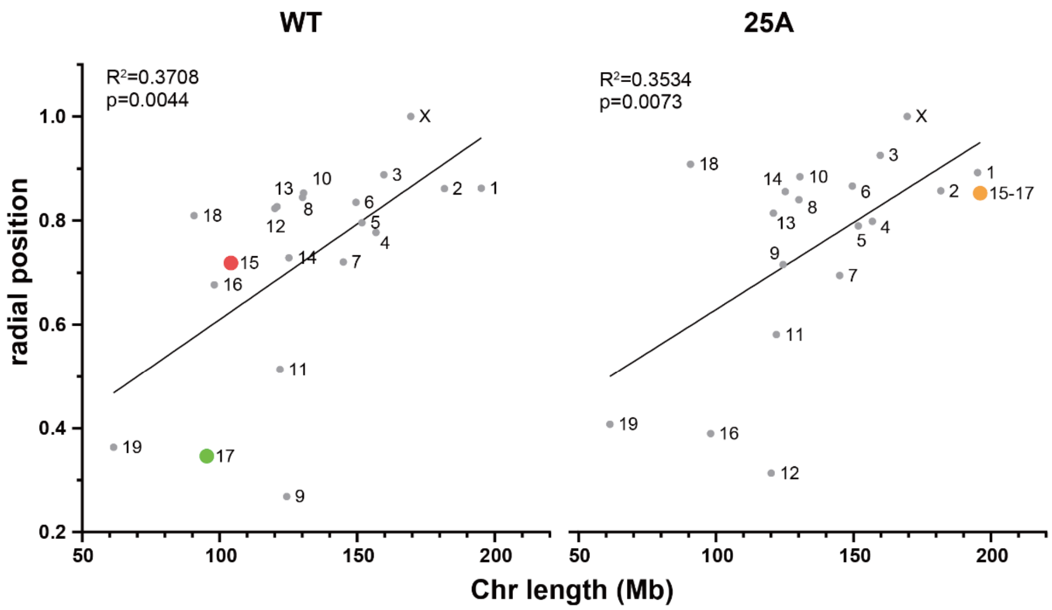

K

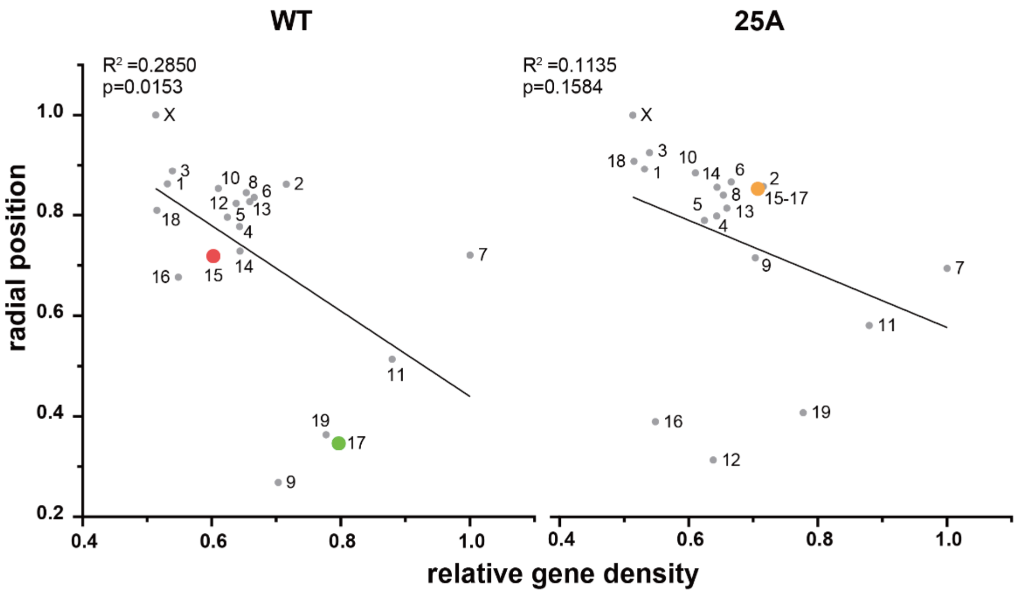

**Figure S2 (related to Figure 2). Chromosomal interactions and territories are changed in 25A cells.**

(A and B) Whole-genome contact matrixes at the 200 kb resolution from WT (A) and 25A (B) cells. (C and D) Circos plots of inter-chromosome interactions among all the chromosomes in WT (C) and 25A (D) cells. Gene densities and contact frequencies were shown as two tracks. Inter-chromosome interactions were colored by types: interactions between Chr15 and Chr17 in red, top 1000 inter-chromosome interactions in blue, and others in grey.

(E and F) Circos plots of filtered inter-chromosome interactions of Chr15 and Chr17 in WT (E) and 25A (F) cells. After normalization, inter-chromosomal interactions with contact scores higher than 500 were shown. The sgRNA target sites on Chr15 and Chr17 in 25A were indicated by yellow bars.

(G) Gene density heatmaps of each chromosome in mouse (GRCm39, GenBank: GCA_000001635.9) at a 5 Mb resolution.

(H) Gene density heatmaps of Chr15 and Chr17 in mouse at a 500 kb resolution.

(I) Chromosome length (left panel) and gene density (right panel) of the genome and the artificially fused Chr15-17 (orange) in mouse. Chr15 in red and Chr17 in green. Black ovals represent the centromere regions. Gene density is shown as the number of genes per Mb on the chromosomes and the dash line indicates the average gene density of the whole genome.

(J) The correlation between relative radial position of chromosomes inferred from 3D genome model with chromosome size in WT and 25A cells. The Y axis represents the relative distance to nucleus center, and the X axis represents the chromosome length in Mb. Pearson's correlation coefficient and p-value were shown (two-tailed).

(K) The correlation between relative radial position of chromosomes inferred from 3D genome model with gene density in WT and 25A cells. The Y axis represents the relative distance to the nucleus center, and the X axis represents the relative gene density normalized by gene density of Chr7 (with the highest gene density). Pearson's correlation coefficient and p-value were shown (two-tailed).

### Figure S3

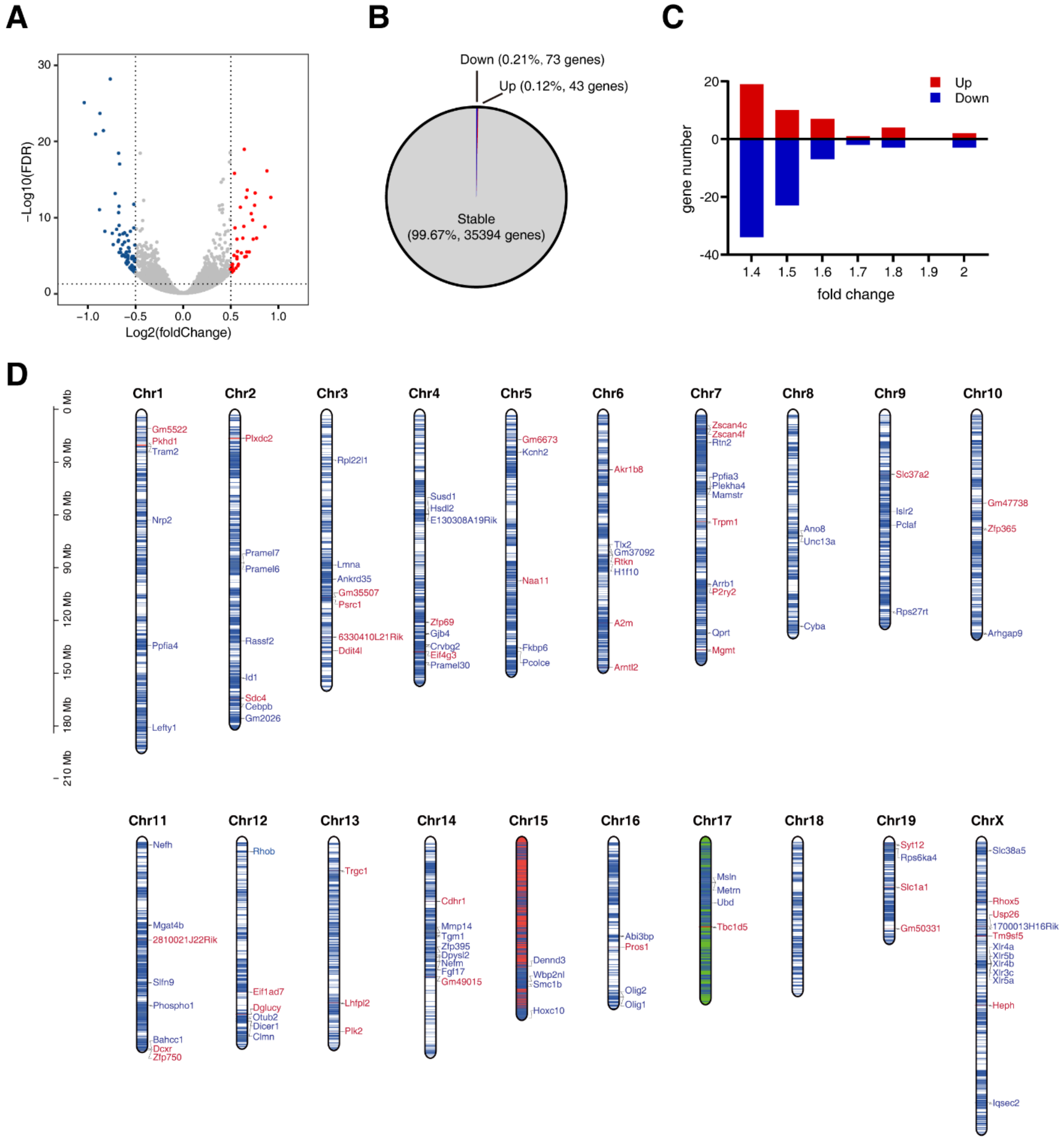

**Figure S3 (related to Figure 3). Transcriptome analyses of 25A cells.**

(A) Volcano plot of differentially expressed genes identified between 25A cells and WT cells. Red dots indicate the upregulated genes and blue dots indicate the downregulated genes (FDR < 5%, log2 fold-change > 0.5 or < -0.5).

(B) Pie graph showing the proportion of up- and down-regulated genes.

(C) Distributions of the number of differentially expressed genes according to the expression fold change. Upregulated genes in red and downregulated genes in blue.

(D) Distributions of differentially expressed genes on each chromosome. The blue lines indicate gene density at a 50 kb resolution. Upregulated genes and downregulated genes are labeled in red and blue on right, respectively.

### Figure S4

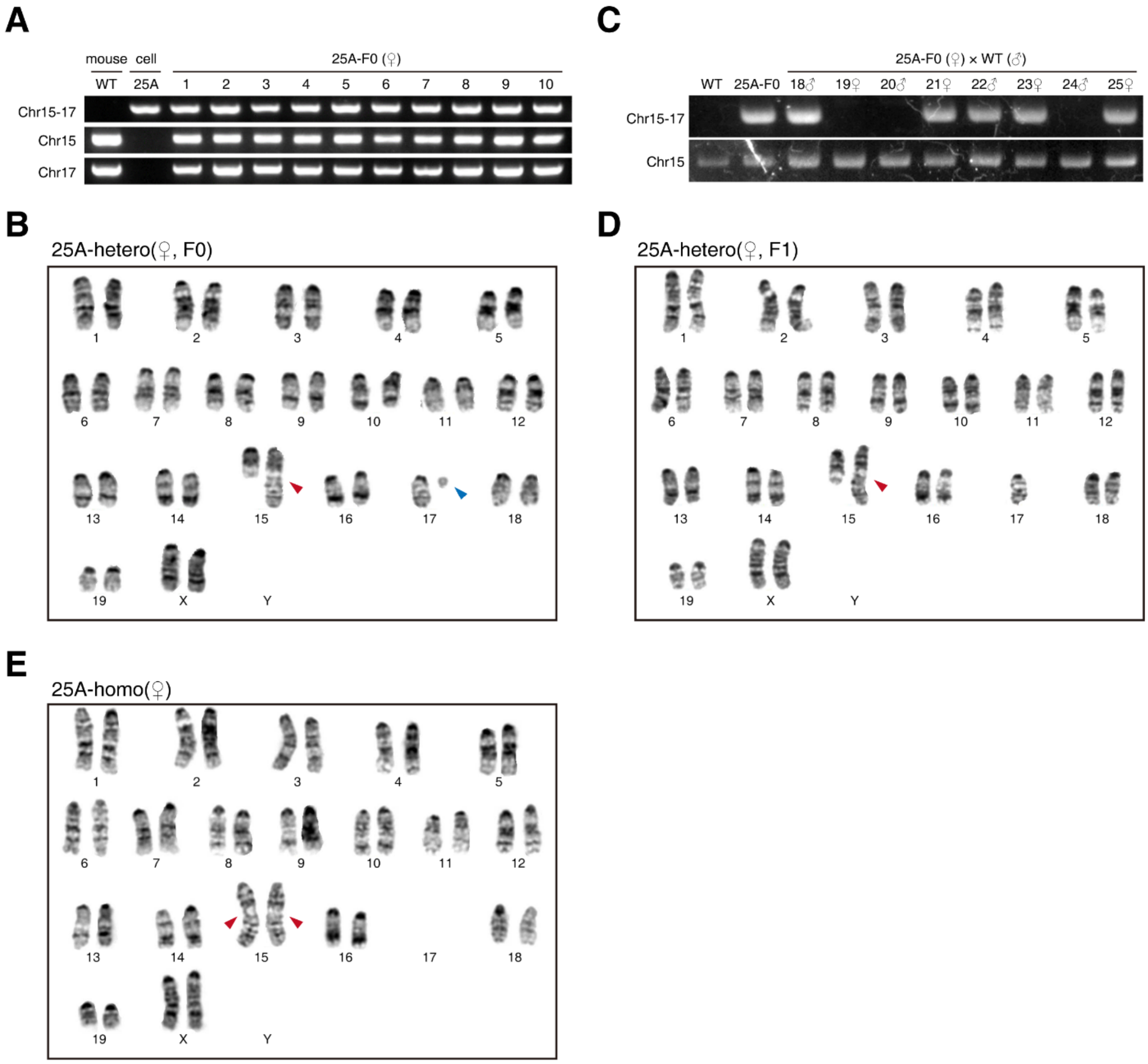

**Figure S4 (related to Figure 4). Validation of heterozygous and homozygous mice containing fusion Chr15-17.**

(A) Genotype analysis of the 25A $F_0$ female mice (heterozygous). Fusion site was detected with primer pairs ‘F1’ and ‘R2’. Original Chr15 and Chr17 was detected with primer pair ‘F1’ and ‘R1’ on Chr15 and ‘F2’ and ‘R2’ on Chr17, respectively. WT mouse and 25A cell line as controls.

(B) G-band karyotype analysis of 25A $F_0$ female mice (38+XX, t(15;17)(F3;A2)). Red arrowhead indicates the fused Chr15-17. Blue arrowhead indicates the mini-chromosome.

(C) Genotype analysis of the 25A $F_1$ mice derived from crossing between 25A $F_0$ female mice and WT male mice. Cross-chromosomal PCR using Chr15-F1 and Chr17-R2 primers for fusion Chr15-17, and inter-chromosomal PCR using Chr15-F1 and R1 for detection of unfused Chr15. WT mouse and 25A $F_0$ female mouse as controls.

(D and E) G-band karyotype analysis of 25A $F_1$ heterozygous female mice (37+XX, t(15;17)(F3;A2)) (D) and 25A homozygous female mouse (36+XX, t(15;17)(F3;A2)×2) (E). Red arrowhead indicates the two copies of fused Chr15-17.

### Figure S5

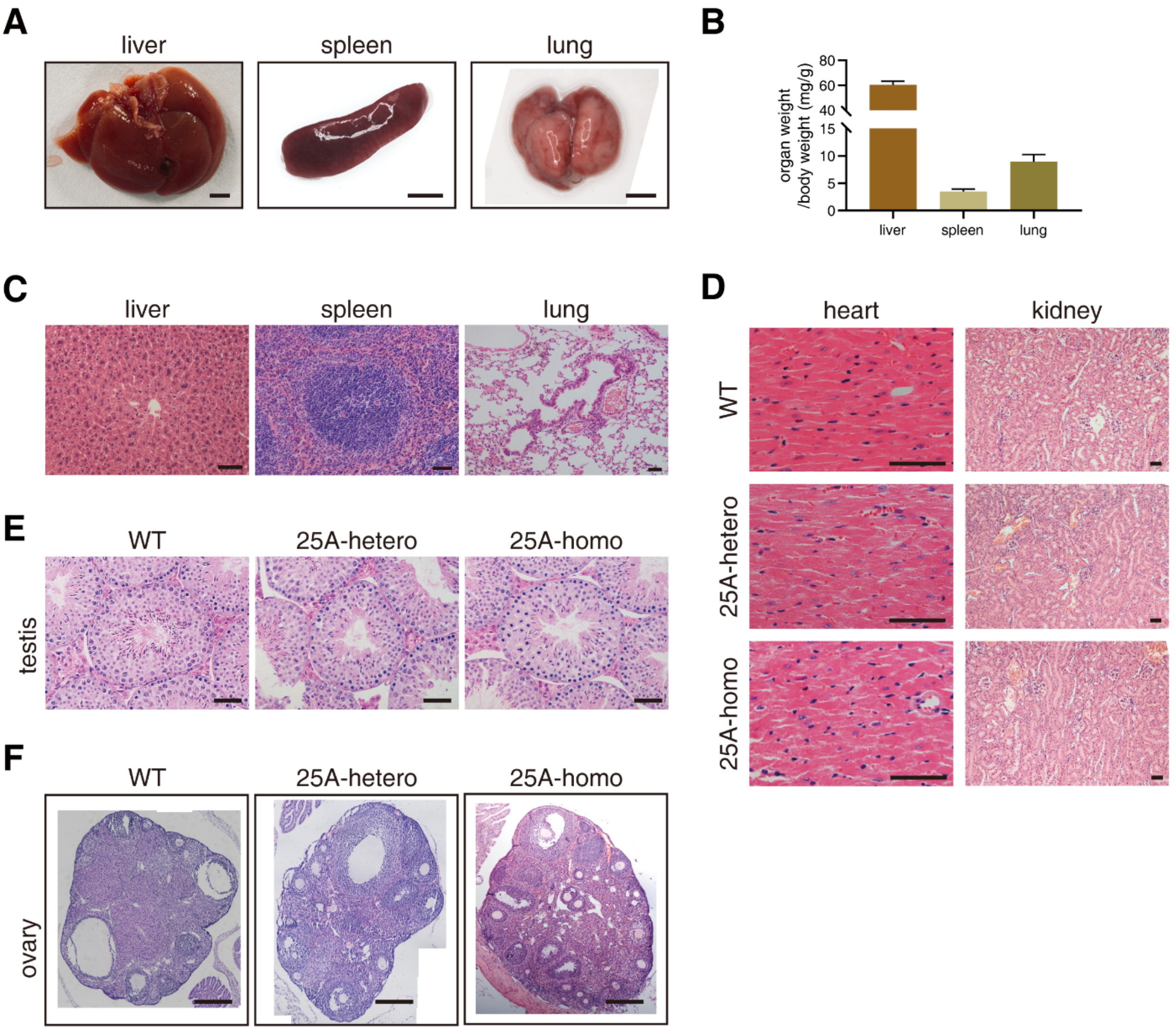

**Figure S5 (related to Figure 5). Phenotypic analyses of 25A mice.**

(A) Morphological features of liver, spleen and lung from 8 weeks 25A (Chr15-17 fusion) heterozygous mice. Scale bar: 300 mm.

(B) Relative organ weight to body weight of liver, spleen and lung of 8 weeks Chr15-17 fusion heterozygous mice. (mean ± sd, two-tailed t-test, n=3).

(C) H&E staining of liver, spleen and lung from 8 weeks Chr15-17 fusion heterozygous mice. Scale bar: 50 μm.

(D) H&E staining of heart and kidney from 8 weeks WT, Chr15-17 fusion heterozygous and homozygous mice. Scale bar: 50 μm.

(E) H&E staining of testis from 8 weeks male WT, Chr15-17 fusion heterozygous and homozygous mice. Scale bar: 50 μm.

(F) H&E staining of ovary from 8 weeks female WT, Chr15-17 fusion heterozygous and homozygous mice. Scale bar: 200 μm.

**Table S1 (related to Figure 5). Complete blood counts of 25A mice.**

| | WT | 25A-hetero | | 25A-homo | |
|---|---|---|---|---|---|
| Item | mean ± sd | mean ± sd | significance vs WT | mean ± sd | significance vs WT |
| WBC(10^9/L) | 5.25±3.38 | 5.83±3.26 | ns (p=0.828843) | 3.78±2.24 | ns (p=0.562918) |
| RBC(10^12/L) | 10.67±0.31 | 11.06±0.69 | ns (p=0.410758) | 10.61±2.42 | ns (p=0.968001) |
| HGB(g/L) | 159.67±5.13 | 167.5±10.02 | ns (p=0.277043) | 157±35.09 | ns (p=0.90265) |
| HCT(%) | 47.3±1.81 | 50.03±3.29 | ns (p=0.257353) | 46.63±9.79 | ns (p=0.91328) |
| MCV(fL) | 44.33±1.78 | 45.25±0.83 | ns (p=0.397388) | 44.1±0.92 | ns (p=0.849783) |
| MCH(pg) | 14.97±0.65 | 15.15±0.17 | ns (p=0.603064) | 14.8±0.1 | ns (p=0.683648) |
| MCHC(g/dL) | 33.77±0.45 | 33.53±0.46 | ns (p=0.517595) | 33.63±0.5 | ns (p=0.749744) |
| PLT(10^9/L) | 994.67±123.79 | 812.25±448.91 | ns (p=0.532498) | 839.33±376.03 | ns (p=0.534046) |
| RDW-SD(fL) | 24.27±1.42 | 22.98±1.23 | ns (p=0.253503) | 22.8±0.26 | ns (p=0.153217) |
| RDW-CV(%) | 14.13±1.15 | 12.93±0.72 | ns (p=0.145445) | 13.63±0.5 | ns (p=0.529544) |
| PDW(fL) | 5.47±0.15 | 5.8±0.37 | ns (p=0.212499) | 5.7±0.26 | ns (p=0.256435) |
| MPV(fL) | 6.93±0.06 | 7.2±0.23 | ns (p=0.11404) | 7±0.1 | ns (p=0.373901) |
| P-LCR(%) | 3.07±0.68 | 4.93±2.11 | ns (p=0.209673) | 2.83±0.42 | ns (p=0.639147) |
| PCT(%) | 0.69±0.08 | 0.58±0.3 | ns (p=0.566542) | 0.58±0.26 | ns (p=0.526606) |
| NRBC#(10^9/L) | 0±0.01 | 0.01±0.01 | ns (p=0.720971) | 0.01±0.01 | ns (p=0.518519) |
| NRBC%(%) | 0.03±0.06 | 0.08±0.1 | ns (p=0.538466) | 0.23±0.25 | ns (p=0.250815) |
| NEUT#(10^9/L) | 0.41±0.21 | 0.47±0.28 | ns (p=0.789945) | 0.49±0.31 | ns (p=0.740705) |
| LYMPH#(10^9/L) | 4.51±3.07 | 5.06±2.81 | ns (p=0.817286) | 3.1±1.86 | ns (p=0.533373) |
| MONO#(10^9/L) | 0.23±0.09 | 0.19±0.1 | ns (p=0.56468) | 0.14±0.05 | ns (p=0.206289) |
| EO#(10^9/L) | 0.09±0.04 | 0.12±0.09 | ns (p=0.692746) | 0.04±0.02 | ns (p=0.116117) |
| BASO#(10^9/L) | 0±0.01 | 0±0.01 | ns (p=0.845671) | 0.01±0.01 | ns (p=0.518519) |
| NEUT%(%) | 8.67±3.71 | 7.98±1.93 | ns (p=0.758016) | 12.6±0.62 | ns (p=0.144183) |
| LYMPH%(%) | 84.37±5.35 | 86.93±2.68 | ns (p=0.437444) | 82±0.56 | ns (p=0.488614) |
| MONO%(%) | 4.87±1.17 | 3.2±0.35 | * (p=0.039024) | 4.07±1.04 | ns (p=0.42574) |
| EO%(%) | 2±0.87 | 1.85±0.48 | ns (p=0.778587) | 1.1±0.1 | ns (p=0.148275) |
| BASO%(%) | 0.1±0.17 | 0.05±0.1 | ns (p=0.646229) | 0.23±0.25 | ns (p=0.491767) |
| PLT-I(10^9/L) | 994.67±123.79 | 812.25±448.91 | ns (p=0.532498) | 839.33±376.03 | ns (p=0.534046) |
| [MicroR(%)] | 9.17±4.75 | 4.88±1.09 | ns (p=0.131909) | 9±1.87 | ns (p=0.957644) |

| | | | | | |
|---|---|---|---|---|---|
| [MacroR(%)] | 2.87±0.12 | 3.08±0.22 | ns (p=0.20372) | 2.93±0.47 | ns (p=0.824042) |
| [TNC(10^9/L)] | 5.25±3.38 | 5.83±3.26 | ns (p=0.828567) | 3.78±2.24 | ns (p=0.564252) |
| WBC-N(10^9/L) | 5.25±3.38 | 5.83±3.26 | ns (p=0.828843) | 3.78±2.24 | ns (p=0.562918) |
| [TNC-N(10^9/L)] | 5.25±3.38 | 5.83±3.26 | ns (p=0.828567) | 3.78±2.24 | ns (p=0.564252) |
| WBC-D(10^9/L) | 5.79±3.62 | 6.06±3.23 | ns (p=0.921042) | 3.84±2.27 | ns (p=0.472849) |
| [TNC-D(10^9/L)] | 5.8±3.62 | 6.07±3.23 | ns (p=0.923465) | 3.84±2.27 | ns (p=0.470984) |
| [NE-SSC(ch)] | 119.1±0.53 | 123.95±4.88 | ns (p=0.155359) | 119.43±0.74 | ns (p=0.559178) |
| [NE-SFL(ch)] | 60.5±0.75 | 61.68±1.88 | ns (p=0.361233) | 61.9±2.38 | ns (p=0.386654) |
| [NE-FSC(ch)] | 69.57±0.78 | 66.5±0.29 | **** (p=0.000703) | 66.37±2.56 | ns (p=0.106832) |
| [LY-X(ch)] | 87.07±0.64 | 85.73±1.11 | ns (p=0.125046) | 86.67±1.31 | ns (p=0.658772) |
| [LY-Y(ch)] | 77.93±5 | 80.4±0.34 | ns (p=0.355477) | 82.13±2.25 | ns (p=0.255285) |
| [LY-Z(ch)] | 60.77±0.76 | 60.1±1.19 | ns (p=0.440313) | 62.13±1.72 | ns (p=0.277168) |
| [MO-X(ch)] | 113±2.7 | 110.55±4.25 | ns (p=0.426794) | 110.87±2.92 | ns (p=0.404843) |
| [MO-Y(ch)] | 154.67±3.66 | 153.08±4.66 | ns (p=0.647619) | 166.27±15.9 | ns (p=0.285511) |
| [MO-Z(ch)] | 71.73±1.54 | 71.7±1.04 | ns (p=0.973789) | 71.37±2.48 | ns (p=0.838469) |
| [NE-WX] | 296.67±28.54 | 267±26.04 | ns (p=0.210718) | 321.33±27.3 | ns (p=0.340194) |
| [NE-WY] | 544.67±43.02 | 547.25±30 | ns (p=0.92836) | 514±97.86 | ns (p=0.64534) |
| [NE-WZ] | 512.33±16.26 | 586.25±23 | ** (p=0.005313) | 579.33±77.69 | ns (p=0.217559) |
| [LY-WX] | 471±27.87 | 460.5±29.85 | ns (p=0.656259) | 473±14.42 | ns (p=0.917428) |
| [LY-WY] | 1065.67±107.45 | 979±90.15 | ns (p=0.29674) | 987.33±77.52 | ns (p=0.36371) |
| [LY-WZ] | 499±14.42 | 478±15.3 | ns (p=0.125347) | 466.33±37.02 | ns (p=0.227502) |
| [MO-WX] | 321±56.96 | 298.5±14.27 | ns (p=0.469702) | 306.33±20.84 | ns (p=0.696834) |

| | | | | | |
|---|---|---|---|---|---|
| [MO-WY] | 303±59.81 | 260.75±85.99 | ns (p=0.502535) | 255.67±120.43 | ns (p=0.574979) |
| [MO-WZ] | 445.33±62.07 | 443±40.46 | ns (p=0.953857) | 349.33±38.84 | ns (p=0.08563) |
| Note: WT, n=3; 25A-hetero, n=4; 25A-homo, n=3 | | | | | |

**Table S2 (related to Figure 1 and Figure S1). CRISPR-Cas9 target sites**

| position | sequence |
|---|---|
| Chr15 (103949158 - 103949177) | GTTCAGGTACATACAACGGT |
| Chr17 (3150411 - 3150430) | GAAAAATTTCCAGATGCCTG |

**Table S3 (related to method). List of primers for PCR and real-time PCR**

| | |
|---|---|
| Chr15-F1 | ATCCAAGTAGCATTAGCTCAGGT |
| Chr15-R1 | GGAGTCTCAGAACCGAGATGTC |
| Chr17-F2 | CTTTTTGAACTCGTATGTAGGCAA |
| Chr17-R2 | GATCAGTTTGGAGACAGAAAAGAG |
| Oct4-F | GGCTTCAGACTTCGCCTCC |
| Oct4-R | AACCTGAGGTCCACAGTATGC |
| Sox2-F | AGGGCTGGGAGAAAGAAGAG |
| Sox2-R | CCGCGATTGTTGTGATTAGT |
| Nanog-F | AAGCAGAAGATGCGGACTGT |
| Nanog-R | ATCTGCTGGAGGCTGAGGTA |
| Pax6-F | TACCAGTGTCTACCAGCCAAT |
| Pax6-R | TGCACGAGTATGAGGAGGTCT |
| Nestin-F | CCCTGAAGTCGAGGAGCTG |
| Nestin-R | CTGCTGCACCTCTAAGCGA |
| KDR-F | GCCCTGCTGTGGTCTCACTAC |
| KDR-R | CAAAGCATTGCCCATTCGAT |
| aSMA-F | GTCCCAGACATCAGGGAGTAA |
| aSMA-R | TCGGATACTTCAGCGTCAGGA |
| PDGFRa-F | GGACTTACCCTGGAGAAGTGAGAA |
| PDGFRa-R | ACACCAGTTTGATGGATGGGA |
| AFP-F | ATCAGTGTCTGCTGGCACGCA |
| AFP-R | GGCTGGGGCATACATGAAGGGG |
| Gata4-F | GGAAGACACCCCAATCTCG |
| Gata4-R | CATGGCCCCACAATTGAC |
| Gata6-F | GGTCTCTACAGCAAGATGAATGG |
| Gata6-R | TGGCACAGGACAGTCCAAG |
| GAPDH-F | AGGTCGGTGTGAACGGATTTG |
| GAPDH-R | TGTAGACCATGTAGTTGAGGTCA |

**Video S1 (related to Figure 2). 3D genome reconstuction of WT.**

**Video S2 (related to Figure 2). 3D genome reconstuction of 25A.**
